# What p53 sees: ATM and ATR activation through crosstalk between DNA damage response pathways

**DOI:** 10.1101/2020.08.04.237131

**Authors:** Elizabeth A. Fedak, Frederick R. Adler, Lisa M. Abegglen, Joshua D. Schiffman

## Abstract

Cells losing the ability to self-regulate in response to damage is a hallmark of cancer. When a cell encounters damage, regulatory pathways estimate the severity of damage and promote repair, cell cycle arrest, or apoptosis. This decision-making process would be remarkable if it were based on the total amount of damage in the cell, but because damage detection pathways vary in the rate and intensity with which they promote pro-apoptotic factors, the cell’s real challenge is to reconcile dissimilar signals. Crosstalk between repair pathways, crosstalk between pro-apoptotic signaling kinases, and signals induced by damage byproducts complicate the process further. The cell’s response to *γ* and UV radiation neatly illustrates this concept. While these forms of radiation produce lesions associated with two different pro-apoptotic signaling kinases, ATM and ATR, recent experiments show that ATM and ATR react to both forms of radiation. To simulate the pro-apoptotic signal induced by *γ* and UV radiation, we construct a mathematical model that includes three modes of crosstalk between ATM and ATR signaling pathways: positive feedback between ATM/ATR and repair proteins, ATM and ATR mutual upregulation, and changes in lesion topology induced by replication stress or repair. We calibrate the model to agree with 21 experimental claims about ATM and ATR crosstalk. We alter the model by adding or removing specific processes, then examine the effects of each process on ATM/ATR crosstalk by recording which claims the altered model violates. Not only is this the first mathematical model of ATM/ATR crosstalk, its implications provide a strong argument for treating pro-apoptotic signaling as a holistic effort rather than attributing it to a single dominant kinase.

## 2 Introduction

All cells receive damage from a wide variety of stressors, each of which contributes to the cell’s ultimate fate. Repair pathways communicate with damage aggregators such as p53, and these aggregators choose whether to promote repair, cell cycle arrest, or apoptosis in response [1, 2, 3]. Without this regulation, the cell can develop tumor-like characteristics such as increased mutation rates and the ability to survive in low-nutrient, low-oxygen, or overcrowded environments [3].

The damage aggregators’ roles would be challenging enough if the signals they received represented the total amount of damage in the cell; however, this is not often the case [3]. Because some signaling pathways only respond to lesions after they undergo preprocessing by the cell, while others activate within minutes after the damage appears, cellular damage induces signals that vary in delay and magnitude [4, 5, 6, 7, 8, 2, 9, 10]. The aggregator must base cell fate decisions on inferences from these signals. Gamma and UV radiation neatly illustrate this concept: both agents cause DNA damage and promote p53-dependent apoptosis, but UV photoproducts weakly activate pro-apoptotic proteins through their repair intermediates while *γ* radiation produces a strong pro-apoptotic response immediately after its induction [11, 12, 13, 14, 15, 8, 16, 17]. Understanding how the cell distinguishes between lethal and non-lethal signals in these two cases is the first step in understanding its decision-making process.

Gamma and UV radiation both produce lesions that bind to p53-activating PIKK-family kinases: ataxia telangiectasia mutated (ATM) binds to the double-strand DNA breaks induced by *γ* radiation, and ataxia telangiectasia mutated and Rad3-related (ATR) responds to repair intermediates of UV photoproducts [4, 5, 6, 7, 8, 2, 9, 10]. ATM and ATR are respectively considered to be the dominant p53-activating kinases, henceforth pro-apoptotic kinases, associated with *γ* and UV radiation [11, 12, 13, 14, 15, 8, 16, 17]. However, recent research shows that *γ* and UV radiation activate both kinases, suggesting that although the cell’s apoptotic decision-making process requires the dominant kinase, it is responding to a pro-apoptotic signal produced by multiple kinases [18, 19, 20, 21]. Several mathematical models explore the apoptotic decision-making response to *γ* or UV radiation based on the dominant kinase activity, but none have attempted to characterize post-radiation signaling as a joint effort between ATM and ATR [22, 23, 24, 25, 26, 27]. We introduce a model of ATM and ATR response to *γ* and UV radiation that incorporates several known or hypothesized modes of crosstalk between the kinases and repair pathways. These include the effects of ATM and ATR on lesion repair, mutual upregulation between ATM and ATR, and changes in kinase binding affinity that occur when the cell alters the structure of a lesion [28, 29, 30, 4, 5]. The model also accounts for changes in ATR activity during S phase and the effects of replication on damage response. From the experimental works we collected, we identified 24 distinct claims about ATM and ATR activation in response to *γ* or UV radiation. We calibrate the model to agree with 21 of these claims, where the final three suggest potential extensions. To explore how each mode of crosstalk affects ATM and ATR signaling, we identify seven key processes and build alternate versions of the model that changes one of these processes. If the alternate model violates one of the original 21 claims, we discuss the implications of this change. Our simulated knockout experiments provide insight on the kinetic effects of different crosstalk mechanisms, and our dose response curves suggest non-dominant kinase activity might be a crucial component of the cell’s response to UV radiation. We intend for this model to be a first step towards studying pro-apoptotic signaling dynamics as a combined effort between multiple damage response pathways.

## 3 Biology background

This section has the following structure: first, we overview properties of ATM and ATR (Section 3.1). We introduce the DNA lesions caused by *γ* and UV radiation in Section 3.2, and review the processes used to repair each lesion in Section 3.3. These repair pathways activate ATM and ATR through the mechanisms laid out in Section 3.4.

### 3.1 ATM and ATR function

Ataxia-telangiectasia mutated (ATM) and ataxia-telangiectasia mutated and Rad3 related (ATR) are two DNA-damage-sensing proteins in the phosphoinositide 3-kinase (PI3K) family (reviewed in [31, 32, 33, 34]). Each has hundreds of substrates, many of which are shared, including p53 and H2A histone family member X (H2AX). ATM and ATR activate p53 by phosphorylating it on Ser15, a required first step toward p53-mediated cell cycle arrest or apoptosis [35]. ATM and ATR also activate checkpoint kinase 2 (Chk2) and checkpoint kinase 1 (Chk1), respectively, two serine/threonine-specific protein kinases that phosphorylate p53 on Ser20 [7, 36, 37, 35]. Ser139 on H2AX is another substrate of both kinases: once phosphorylated, the histone allows damaged DNA to unwind so that repair proteins have easier access to the lesion [7, 36, 37, 35].

While ATM and ATR are similar in structure and function, they are activated by different types of DNA damage. Inactive ATM is a dimeric protein that splits into two monomers when it encounters a double-stranded break. It then phosphorylates itself on Ser1981 and other sites, activating its kinase function, and the monomers bind to opposite sides of the break [4, 5, 6, 7]. ATM has a multitude of substrates, including damage response proteins such as p53 and Mdm2, and DSB signaling proteins such as Mre11, Rad50, Nbs1, H2AX, MDC1, 53BP1, and BRCA1 [7, 36, 37, 35]. The ATM-mediated kinase cascade drives the formation of large clusters of proteins, called foci, around the DSB [26, 38]. ATM is classically associated with DSB-causing agents such as *γ* radiation and doxorubicin [11, 12, 13, 14].

ATR is activated by single-stranded DNA (ssDNA) bound to replication protein A (RPA) (reviewed in [33, 34, 39, 40]). ATR must be bound to ATR-interacting protein (ATRIP) to associate with RPA-bound ssDNA and must interact with other proteins, including topoisomerase 2-binding protein 1 (TopBP1), to become active [30, 34]. Active ATR is autophosphorylated on multiple sites, including Ser428, Ser435, and Thr1989 [41]. Rather than binding directly to lesions, ATR interacts with the strands of RPA-bound ssDNA created as repair intermediates or through cellular misprocessing [9, 34].

ATR promotes G2 arrest and is primarily active during S phase. That is, the cell can create RPA-bound ssDNA during all stages of the cell cycle, but ATR reacts more strongly to ssDNA created during replication stress [42]. Cell-cycle-specific regulators such as XPA, an important cofactor, promote ATR binding during S phase [43]. During G1 phase, XPA is mostly sequestered in the cytosol, and in G2, XPA is mostly sequestered in the nucleus [43]. At the start of S phase, XPA is initially cytosolic, but imports to the nucleus in a p53-dependent manner after UV damage [43]. This creates a self-regulatory positive feedback loop for ATR. Active ATR also upregulates RPA, another mechanism of positive feedback [44]. ATR is classically associated with lesions that are excised during repair, such as those produced by UV radiation and cisplatin [15, 8, 16, 17].

### 3.2 DNA-Damaging Agents

The two damaging agents we consider in this work are *γ* and UV radiation. Below, we outline the DNA lesions they induce and their relationship with ATM and ATR.

#### 3.2.1 *γ* radiation induces single-strand breaks (SSBs), double-strand breaks (DSBs) abasic sites, oxidized bases, and multiply damaged sites (MDS); is associated with ATM

Exposing a cell to *γ* radiation, a low Linear Energy Transfer (LET) form of ionizing radiation, creates several types of DNA lesion, including single-strand breaks (SSBs), sites where the base dissociates while leaving the sugar backbone is intact (abasic sites, AP), and oxidized base products (reviewed in [45, 46]). Gamma radiation releases multiple ions to release around sites where it passes through DNA, creating dense clusters of lesions. The cell easily repairs isolated lesions, but clustered lesions are often persistent and harder to repair [46].

Clustered lesions, also called multiply damaged sites (MDS), vary in complexity from two lesions to three or more [46]. The most well-known clustered lesion is the double-strand break (DSB), which is produced when complementary base pairs dissociate between two close-by SSBs, which must be on opposite strands and separated by ten or fewer base pairs [47, 48]. The cell does not use the same pathways to repair clustered lesions as it does to repair their constituent components [49, 50]. Eukaryotic cells have four pathways to repair double-strand breaks; multiply damaged sites undergo unpredictable configuration-specific repair processes [49].

A 1988 meta-study measured four complex DSBs, 36 simple DSBs, and 400 MDS induced for every 1 Gy of *γ* radiation [45]. In the range of 0–50 Gy, the number of lesions increases linearly with the dose [45].

ATM becomes robustly activated less than 2 minutes after exposure to *γ* radiation [51, 52]. Within 15 minutes, it activates p53 as well [22]. We observe this rapid activation even in response to low doses of *γ* radiation, suggesting that a fast signal does not necessarily imply a lethal amount of damage [31]. Instead, the p53 pathway appears to distinguish between *γ* radiation doses by the signal duration, where longer-lasting signals become lethal [24].

#### 3.2.2 UV radiation induces thousands of photoproducts; is associated with ATR

UV radiation induces lesions on DNA collectively known as UV photoproducts [53, 54]. UV-induced free radicals react with pairs of pyrimidines to create bulky adducts, mostly cyclobutane pyrimidine dimers (CPDs) or pyrimidine (6–4) pyrimidone photoproducts (6–4PPs) [53, 54]. Both bulky lesions stall high-fidelity DNA and RNA polymerases [54, 55, 56]. The majority of these photoproducts are CPDs (~75% of UV photoproducts), which take five times longer to repair than 6–4PPs (~25%) [57, 54, 58].

UV radiation induces approximately 4500 photoproducts for each 1 J/m^2^ [59]. These photoproducts do not cluster and are roughly Poisson distributed across the genome [60]. Exposed keratinocytes can have 10^5^ photoproducts after a day in the sun [61].

The UV damage response is cell-cycle specific: ATM and ATR respond to photoproducts in S and G2 phase, but not G1 [62]. Even during S phase, ATR-phosphorylated Chk1 and *γ*H2AX do not appear immediately after UV damage, although CPDs are present, and their concentrations continue to slowly increase in normal human fibroblasts after 20 hours [62]. The number of ssDNA strands created during S phase also depends on whether the cell was actively replicating when it was irradiated. If the cell is in G1/G2 phase, some photoproducts will be repaired by TC-NER and GG-NER before S phase starts; this would not happen for damage induced during S phase. In addition, Li et al. induce significantly more CPDs in HeLa cells by exposing them to UV during S phase instead of G1, possibly due to changes in chromatin configuration or increased amounts of DNA [43].

### 3.3 Classes of lesion

Gamma radiation induces single-strand breaks, abasic sites, oxidized bases, multiply damaged sites, and double-strand breaks. UV radiation induces UV photoproducts. In this section, we describe how the cell repairs this damage.

#### 3.3.1 Single-strand breaks (SSBs), abasic sites, and oxidized bases are repaired by break excision repair (BER)

A single-stranded region of DNA with missing bases and a broken sugar backbone is called a single-strand break (SSB) [45]. The cell encounters up to 10^4^ SSBs per day, some of which it creates through endogenous processes such as DNA unwinding [63, 64, 61]. Although these breaks are ubiquitous in wild-type cells and do not promote apoptosis on their own, they can become dangerous if left unrepaired. The DSBs created by ionizing radiation, for example, are two SSBs within ten base pairs of one another [65]. SSBs can also induce DSBs through replication stress: when a replication fork separates a broken strand of DNA from its complement, the break in the template strand and its newly synthesized complement form a single-ended DSB [66].

The global single-strand break repair process, active during all stages of the cell cycle, begins when a PARP-family protein, frequently PARP-1, attaches to the SSB (reviewed in [67]). The PARP-family protein attaches long chains of poly(ADP-ribose) (pADPr) to itself and other proteins and recruits other repair proteins to the damaged site [67]. One notable early responder to pADPr chains is X-ray repair cross-complementing protein 1 (XRCC1), a scaffolding protein that is also capable of stimulating the activity of downstream enzymes [67, 68]. XRCC1 and pADPr chains both attach to SSBs less than two minutes after irradiation, but while pADPr chains dissociate from the break within 15 minutes, XRCC1 foci persist after 30 minutes [68].

There are two pathways the cell can use for gap filling and DNA ligation, depending on whether the resected region requires a short (1 nt) or long (2-12 nt) nucleotide patch [67]. To create a short patch, XRCC1 recruits DNA polymerase *β* (Pol *β*) and DNA ligase III*α*, which respectively synthesize the missing base and reconstruct the DNA backbone [67]. This short-patch repair process can resynthesize most endogenous SSBs [67]. If the SSB has damaged 3’ and 5’ boundaries, the cell must excise the damaged ends and replace the damaged region with a longer nucleotide patch [67]. DNA ligase I aids in synthesizing longer nucleotide chains by joining a PCNA loading ring and Pol *δ*/*E* to the sugar backbone [67]. Excising the damaged bases creates a short region of ssDNA, but since the SSB repair pathways do not involve RPA, they do not promote ATR [67].

Two additional types of lesion induced by *γ* radiation, abasic sites and oxidized bases, undergo the same repair process as SSBs after being excised from the DNA (reviewed in [69]). An additional pathway repairs the replication-stress byproducts created by SSBs [70]. Because the replication-stress byproducts of SSBs are DSBs, we address this repair process in the DSB section [70].

#### 3.3.2 Multiply damaged site (MDS) repair can produce double-strand breaks (DSBs)

Repairing clustered lesions is not as straightforward as addressing each simple lesion individually: it depends on the configuration of each lesion cluster (reviewed in [50]). For an MDS composed of abasic sites, single-strand breaks, and oxidized bases, each individual lesion in the cluster would be repaired by break excision repair (BER) [50]. Abasic sites and oxidized bases must be excised during the first step of BER—but if the excised site is close to a single-strand break on the opposite strand, the complementary strands can dissociate to form a DSB [71, 50]. Blaisdell et al. provide a detailed of which lesion configurations and enzyme interactions lead to DSB formation [71]. ATM does not interact with MDS, but responds to the DSBs created as repair intermediates [4, 5, 6, 7].

#### 3.3.3 Double-strand breaks (DSBs) are repaired by non-homologous end joining NHEJ), alternate end joining (AEJ), single-strand annealing (SSA), or homologous recombination (HR), depending on the length of their single-strand overhangs

Double-strand breaks (DSBs) lead to aneuploidy and deletion mutations, which threaten the cell’s survival [14, 13]. Eukaryotic cells prevent these genome modifications by leashing together the loose ends of a DSB before they degrade or move away from each other, and, if this method fails, promoting a high-fidelity form of DSB repair [72, 14]. The first method relies on fast detection; joining two DSB ends quickly after their induction increases the chances that they originated from the same strand [4]. Fast DSB detection occurs in a variety of vertebrate cell lines, where protein tethers activate within 2 minutes after irradiation [51, 52]. DSBs that include additional simple lesions beyond the two SSBs, or that are buried in the chromatin structure, may take longer to repair. Two of the processes used by the cell to repair these persistent DSBs are highly mutable [73]. Eukaryotic cells do have a high-fidelity DSB repair process, which is mostly active during S phase [14]. Because some DSBs are challenging to repair but leaving them unjoined can lead to lethal consequences, DSB repair is a complicated multistep process.

There are four DSB repair pathways: classical non-homologous end joining (NHEJ), alternate end joining (AEJ, also known as alt-EJ or B-NHEJ), single-strand annealing (SSA), and homologous recombination (HR) [73, 72, 14]. Human cells primarily repair DSBs by NHEJ, a fast process that requires the fewest steps [73, 72, 14]. When a DSB persists through NHEJ, the next-best option is HR, a slow but error-free pathway that mainly operates during S phase [73, 72, 14]. AEJ and SSA are less common and highly mutagenic [73]. Several factors influence the cell’s choice of repair pathway, including the cell cycle, break complexity, chromatin structure around the break, and configuration of base pairs around the break (reviewed in [72, 50]). We briefly introduce each repair pathway.

Non-homologous end joining is the predominant DSB repair pathway in vertebrates (reviewed in [14]). It is most frequently initiated when Ku, a ringlike protein composed of subunits Ku70 and Ku80, threads itself onto broken ends of DNA[74, 75]. Ku preserves the structure of the DSB by guarding against accidental breakage or extensive end resection, like the aglets on a shoelace [74, 75]. It then attracts XRCC4-DNA ligase IV complex, the NHEJ ligase that joins the loose ends of DSBs [76, 77, 78]. This complex can only connect single-stranded overhangs if they have some degree of microhomology, typically 2-4 bp [14]. If the overhangs are incompatible—e.g., if the DSB includes three or more lesions—they may undergo multiple rounds of nucleation and DNA polymerization before XRCC4-DNA ligase IV can process them (reviewed in [73]). Ku promotes nucleation by activating DNA-PKcs, a phosphoinositide 3-kinase (PI3K), which attracts Artemis, a nuclease that processes the overhangs [79, 80].

Following *γ* irradiation, NHEJ repairs half of the radiation-induced DSBs within 20 minutes and 80% or more within 2 hours. The remaining DSBs typically undergo further processing by the cell before repair and can persist for days [50]. The contrast in speed between NHEJ and other DSB repair pathways gives DSB repair kinetics a characteristically biphasic structure [81, 13, 82]. NHEJ proteins compete with endonucleases that cleave the ends of a break to produce longer ssDNA overhangs, a process called end resection (reviewed in [73]). Once a break undergoes end resection, it must be repaired by AEJ, SSA, or HR, because NHEJ can only join single-strand DNA (ssDNA) tails that are 0-10 base pairs long [73]. In humans, this end resection is initiated by a complex of meiotic recombination 11 (Mre11), Rad50, and Nijmegen breakage syndrome 1 (Nbs1), referred to as the MRN complex (reviewed in [4, 83]). End resection begins when CtIP-activated MRN complex cleaves the DSB ends to create 15-100 bp ssDNA overhangs on either side of the break [4, 83]. Lengthening the tails may expose longer homologous regions, allowing the cell to repair the break using either AEJ or SSA, where AEJ requires regions of microhomology (4-6 bps) and SSA requires at least 20 matching nucleotide pairs (reviewed in [73, 72]). For AEJ, once the ssDNA tails bind across a region of microhomology, Pol *θ* stabilizes the short homologous region and fills the gaps on flanking ssDNA [72]. DNA ligase I or III then joins the newly synthesized strands to the existing dsDNA. To expose the longer homologous regions required for SSA, the ssDNA tails undergo multiple rounds of end resection [72]. Several helicases and nucleases can mediate this resection, including DNA2, BLM, WRN, CtIP, and EXO1 [72]. Replication protein A (RPA) binds to and stabilizes the newly exposed ssDNA [72]. Once homologously bound, the dsDNA regions have unpaired nucleotide tails at one end and ssDNA on the other. Nuclease complex XPF-ERCC1, composed of xeroderma pigmentosum F and excision repair cross-complementation group 1, cleaves these loose ends [72]. Lastly, Rad52 helps anneal the homologous ssDNA pairing and polymerases/ligases repair the ssDNA gaps [72]. SSA can delete large regions of the genome and AEJ can introduce short insertions or deletions depending on local homology; as a result, these repair pathways are highly mutagenic [72].

In HR, ssDNA tails are matched with an intact sequence on the homologous chromosome or sister chromatid (reviewed in [14]). The cell can then use the undamaged sequence as a template for repair. Out of all four repair processes, HR requires the longest ssDNA tails [84, 85]. Rad51 initiates the extensive end-resection needed for HR, then, like Rad50, tethers the overhangs to their homologous template strand through the “molecular Velcro” mechanism [86]. The homologous template region exchanges strands with the ssDNA tail and DNA Pol I synthesizes the complement of the exposed ssDNA [14]. Detangling the newly created dsDNA involves several DNA processing enzymes whose properties are outside the scope of this work.

A subclass of HR, break-induced repair or BIR, repairs DSBs that are created as byproducts of replication stress (reviewed in [70]). When a replication fork separates a single-strand break from its complement strand, the broken strand and newly synthesized complement form a DSB with only one end [70]. These single-ended DSBs cannot be repaired by end-joining processes. Instead, after the ssDNA overhang has been end-resected and bound by Rad51, it invades the complementary dsDNA on the homologous chromosome [70]. DNA polymerases bind to ssDNA on both sides and the two template strands become a new replication fork [70]. Rad51-dependent BIR is a slow process: it does not initiate until 2-4 hours after a single-sided DSB forms, and it takes an additional 30 minutes to resynthesize the replication fork [87].

Minutes after a DSB appears, mediators of all four repair pathways compete at the break site to determine its ultimate fate [88, 72]. Binding interactions, break location, and timing all influence the choice of repair pathway [88, 72]. The first competitive interaction is between Ku and MRN complex: Ku shields the short ssDNA tails from being nucleated and MRN complex initiates end resection [72]. If Ku persists long enough for NHEJ to repair the break, the process ends. Otherwise, if NHEJ is too slow, MRN complex irreversibly starts strand cleavage, so that one of the other three pathways must repair the break [72]. NHEJ is prevalent because Ku is highly abundant in human cells: Beck et al. measured 100 times more Ku subunits than MRN complex subunits in a quantitative analysis of protein concentrations in U2OS (human osteosarcoma) cells [73, 89]. Thus, the longer a DSB is left unrepaired, the higher the chance it will be repaired by one of the three other pathways.

Competitive binding between RPA and Rad51 determines whether HR or one of the other two processes repairs the break [72]. We focus on three important Rad51 regulators: replication protein A (RPA), breast cancer type 2 susceptibility protein (BRCA2), and breast cancer type 1 susceptibility protein (BRCA1) [86]. RPA coats the break’s ssDNA tails during NHEJ and the first rounds of end resection, so that Rad51 cannot initiate HR without first displacing RPA [86, 90]. BRCA2 increases the chances of Rad51 outcompeting RPA: it prevents ssDNA-bound Rad51 from degrading and stabilizes Rad51 microfilaments [86, 90]. BRCA1 interacts with both the MRN complex and Rad51 to promote end resection [72]. Many proteins that promote Rad51 activity, including BRCA1 and BRCA2, are most active during S and G2 phases [72]. As a result, HR is more common during replication than during other stages of the cell cycle [91]. Because the amount of DNA in the cell doubles during replication, HR proteins can also locate homologous DNA more easily in S and G1 phase.

Multiple signaling factors, including BRCA1, MRN complex, ATM, and *γ*-H2AX, are involved in all four repair pathways (reviewed in [4, 83]). ATM and MRN complex alter the break site within minutes: MRN complex aggregates proteins and bunches DNA around the break, and ATM phosphorylates H2AX to Ser139H2AX/*γ*H2AX to unwind the damaged site from the chromatin structure [4, 5]. This reconfiguration attracts proteins that creates a bulky mass, called a ‘focus,’ around the DSB [4, 5]. Foci can contain multiple break sites and become larger as the DSB persists.

#### 3.3.4 UV photoproducts are repaired by nucleotide excision repair (NER)

UV radiation interacts with DNA to create bulky lesions called photoproducts [53, 92, 54, 44]. These lesions become mutagenic during replication, where they are too bulky to be processed by high-fidelity DNA or RNA polymerases and must be processed by specialized, less accurate polymerases [92, 54, 44]. UV photoproducts can be detected in one of three ways: by stalling RNA polymerase during translation, stalling DNA polymerase during replication, or being flagged by regulatory proteins [93]. Photoproducts on actively-transcribed genes or stalled replication forks are recognized faster than those found by DNA-proofreading proteins [21, 94, 93]. Thus, the rate at which a single photoproduct is detected depends on its position in the genome and the cell’s stage in the cell cycle, making it difficult for the cell to infer the extent of radiation damage.

Of the two primary types of photoproduct, CPDs and 6–4PPs, we use CPDs as the representative photoproduct because they are more persistent than 6–4PPs and three times as common [57, 54, 58]. The cell uses two different nucleotide excision repair pathways to identify CPDs: transcription-coupled nucleotide excision repair (TC-NER) for those on actively transcribed regions and global genome nucleotide excision repair (GG-NER) in all other parts of the genome [93, 58, 95]. A third pathway, trans-lesion synthesis (TLS), fills in photoproduct-induced gaps in replicated DNA [93].

Transcription-coupled NER repairs lesions on actively transcribed genes. When RNA polymerase II (RNAP II), the enzyme responsible for creating almost all cellular mRNA, encounters a photoproduct, it halts DNA translation and stalls at the damaged cite [96]. Helper protein Cockayne syndrome B (CSB) stabilizes the damaged site through its ATP-dependent chromatin remodeling ability, attracting helper protein Cockayne Syndrome A (CSA) [97, 98]. CSA then forms a complex with DDB1-CUL4A-ROC1 that functions as an E3 ubiquitin ligase [97, 99, 98, 96]. This complex ubiquitinates RNAP II, prompting other factors to hyperphosphorylate its largest subunit [96]. RNAP II then dissociates from the damaged strand, clearing space on the DNA for downstream NER factors to bind [96, 93].

The stalled transcription apparatus attracts transcription factor IIH (TFIIH) complex, which unwinds the damaged DNA and facilitates NER complex binding [100, 93]. XPA and RPA define excision sites on the remaining strand [101]. RPA is polar, which may help guide 3’ or 5’ cleavage enzymes to the correct region [101]. XPA, TFIIH, and XPC cooperate to prevent NER machinery from incorrectly incising undamaged DNA, a phenomenon Luijsterberg et al. name “kinetic proofreading” [102]. Endonucleases must interact with all three proteins to cleave the DNA, and because both XPC and XPA bind weakly, this most likely happens when the DNA strand is improperly bent [102]. Once they detect all three proteins, nucleases XPG and XPF/ERCC1 (excision repair cross-complementation group 1) cleave the 3’ and 5’ ends of the damaged site, respectively [100, 93]. Proliferating-cell nuclear antigen (PCNA) recruits Pol*δ*, *ε*, or *κ* to resynthesize the excised region [100, 93]. The NER proteins may not bind in this order, but the damaged strand always undergoes the same conformational changes during repair [102].

Global genome NER handles photoproducts on-transcribed regions. A complex of DDB2, DDB1, Cul4A, and Roc1, the analog of the DDB1-CSA E3 ubiquitin ligase involved in TC-NER, binds to damaged chromatin [103, 99]. After exposure to UV radiation, this complex releases CSB, a ubiquitin ligase suppressor that targets DDB1-CSA, XPC, and DDB2 [99, 103]. A complex composed of XPC and HR23B attracts TFIIH, after which GG-NER proceeds identically to TC-NER [104, 93].

#### 3.3.5 Stalled replication forks are repaired by trans-lesion synthesis (TLS)

If left unrepaired before the start of S phase, lesions can physically block DNA synthesis proteins from moving forward on a template strand [105, 106]. The resulting damaged site is called a stalled replication fork. When this blockage occurs, the lesion is bypassed and DNA replication machinery may re-prime at a downstream site [106]. Neither TC-NER nor GG-NER can repair lesions on single-stranded DNA, since they rely on excising the damaged region and using the complementary strand as a template; rather, the gaps created by this process are filled by a third repair pathway known as trans-lesion synthesis (TLS) [105, 106]. Photoproducts on unprocessed DNA can still be repaired by TC-NER or GG-NER [93].

High-fidelity DNA polymerases cannot synthesize base pairs across from bulky adducts such as photoproducts. Instead, translesion synthesis polymerases (Pols. *β*, *ζ*, *η*, *θ*, *ι*, *κ*, *λ*, *µ*, *ν*) pair complementary nucleotides to these lesions [107]. Each polymerase can repair some lesions with high fidelity (e.g., Pol *η* and TT-CPDs) but mis-repair others (e.g., Pol *η* and 6–4PPs), such that TLS polymerases are specialized to address a wide geometry of DNA lesions [107]. When PCNA encounters one of these adducts, it undergoes post-translational modifications to recruit these alternate polymerases to the damaged site: for example, when Rad6-Rad18 complex ubiquitinates PCNA, the altered loading ring has a higher affinity for Pol *η* [108]. After one polymerase processes the bulky adduct, a second TLS polymerase can begin synthesizing DNA across from the single-stranded RPA-bound gap between the lesion and the re-primed site [109]. These two operations are not necessarily performed by the same TLS polymerase [107]. This process restores the complementary strand but keeps the original photoproduct intact. NER must repair the original lesion after TLS is complete.

### 3.4 Interactions between repair pathways, ATM, and ATR

Having introduced each of the repair pathways, we now address how they interact with ATM and ATR. The DSBs caused by *γ* radiation directly bind to ATM and ATR, where end resection increases the amount of single-stranded DNA available for ATR binding [110]. The single-stranded DNA regions created as repair intermediates for UV photoproducts activate ATR but can also break to form DSBs [111]. Moreover, ATR can directly phosphorylate ATM, but ATR lacks the proper binding motif [28]. ATM and ATR can influence repair pathways once activated: they promote DNA repair through structural modifications to DNA and also suppress cell cycle progression [44, 112]. These interactions form the foundation for holistically understanding ATM and ATR activation dynamics after exposure to both forms of radiation.

#### 3.4.1 DSBs upregulate ATM and ATR

ATM is essential for later stages of DSB repair. Even though foci containing *γ*H2AX and active ATM form within 2 minutes of irradiation, inhibiting ATM does not delay DSB repair until after 10 minutes [88, 51, 52, 113]. In ATM knockout cells (A-T cells), between 10–25% of DSBs took longer to repair [114, 115]. ATM therefore appears to most strongly influence repair for a small number of persistent DSBs. Several factors influence the speed of DSB repair, including the number of simple lesions clustered around the break and the surrounding chromatin structure (reviewed in [114, 49]). Breaks buried in the chromatin structure, in particular, rely on ATM’s histone-modifying properties to reveal it to repair proteins [13, 113, 91]. Because ATM interacts with persistent breaks, it produces a prolonged signal in response to *γ* radiation.

Because most complex breaks undergo end resection, we might expect ATM to interact with end-resected breaks—but once a DSB undergoes end resection, it loses the blunt ends that interact with ATM [116]. DSBs with extended single-stranded overhangs do not bind to ATM in environments where they are isolated from all other regulatory proteins [116]. DSB foci could prolong ATM activation by clustering ATM-phosphorylating proteins, such as ATM itself, around the break, such that blunt ends are only needed to initiate focus formation [4, 5].

ATR binds to the RPA-coated ssDNA strands on end-resected DSBs. Active ATR promotes HR and other forms of complex DSB repair (reviewed in [84, 50]). But ATR foci can form around DSBs even before end resection: in cells with knocked-out CtIP, a factor required to initiate end resection, ATR still accumulated around DSB sites [117]. The ATR signal produced in CtIP-knockout cells is transient, so the authors conclude end resection is required for sustained ATR checkpoint signaling [117]. To become active before detectable end resection, ATR may interact with the short ssDNA tails on non-end-resected DSBs [117]. Lastly, visible ATR foci form around DSBs in G1 phase cells, even though ATR targets do not respond to UV damage in G1 phase [20]. It is therefore possible for ATR to become active outside of S phase.

We may question whether ATR binds to extensively end-resected DSBs since this form of end resection requires Rad51 to outcompete RPA on the ssDNA tails. Experimentalists have observed that even if RPA is temporarily displaced, it can rebind to Rad51-coated ssDNA [90]. The two proteins are known to bind to each other and frequently colocalize [90]. Knowing this, we assume DSBs in the extensively end-resected class are still capable of activating ATR.

#### 3.4.2 UV photoproducts and stalled replication forks upregulate ATR and ATM

ATR is the PIKK-family kinase associated with UV damage because it binds to the RPA-bound ssDNA strands created when photoproducts are excised or cause replication forks to stall [109, 62, 118, 16]. This delays the rate at which ATR reacts to UV damage: not only can UV photo-products evade detection for hours, ATR binds to repair intermediates rather than photoproducts themselves [109, 62, 118, 16]. Contrast this with ATM, which locates DSBs within a minute of irradiation [51, 52]. ATR activates several proteins involved in UV repair, including XPA, H2AX, and Pol*η*. It regulates XPA through two positive feedback loops: it increases the binding affinity of XPA to ssDNA by phosphorylating it on Ser196 and increases the nucleic import of XPA during S phase through a p53-dependent mechanism [44, 43]. ATR modifies histones using the same mechanism as ATM, phosphorylating H2AX to *γ*H2AX [119]. Lastly, ATR phosphorylates Pol*η* on Ser601, which promotes its polymerase activity during S phase [120].

The cell detects photoproducts on transcribed strands more effectively than photoproducts in less active regions of the genome. Within 4 hours of irradiating fibroblasts with UV, 68% of all CPD repair byproducts are located on transcribed strands, suggesting TC-NER performs most early CPD repair [92]. CPDs detected by GG-NER can persist more than 36 hours after UV exposure, but CPDs on transcribed strands are repaired within ~10 hours [58, 95, 92, 21]. We assume this difference is caused by TC-NER being able to detect CPDs more efficiently since the downstream repair pathway is the same for both processes. We also expect photoproducts to persist longer during S phase since replication-stress-inducing photoproducts must undergo both TLS and NER to be repaired [44]. Because TLS does not need to be completed by the end of S phase, photoproducts induced during S phase can persist into G2 phase [107, 106, 62].

ATR suppresses origin firing near sites of DNA damage [121]. That is, with fewer replication forks in the cell, fewer replication forks stall on lesions, preventing new RPA-bound ssDNA sites from forming. Hence, ATR can affect the rate at which its associated lesion sites appear by hindering replication in S and G2 phases.

UV photoproducts do not directly interact with ATM. However, the ssDNA created during photoproduct repair has a low chance of breaking during NER to form a DSB [111]. We refer to this process as endemic breakage.

#### 3.4.3 ATR upregulates ATM

Although ATM and ATR appear to compartmentalize two different classes of damage, researchers have discovered situations where ATM acts downstream of ATR, ATR acts downstream of ATM, and knocking down one of these kinases drastically changes the activity of the other [122, 18, 123, 19, 21]. Hannan et al. (2002) reported that cells with defective ATM (A-T cells) were slower in repairing UV damage, and showed higher concentrations of p53 4 hours after UV induction than wild-type cells [122]. This suggests that ATM plays a nontrivial role in UV damage signaling despite being unable to bind to photoproducts or their cellular processing intermediates. This result was further confirmed by Stiff et al. (2006), who also found that silencing ATR suppresses the ATM response to UV radiation and hydroxyurea [19]. Earlier that year, Jazayeri et al. (2006) showed that ATR was activated during replication as part of the cell’s response to *γ* radiation and that ATR activation after *γ* irradiation required ATR [18, 123]. Jazayeri et al. also found that A-T cells had significantly lower concentrations of RPA-bound ssDNA after *γ* radiation exposure and identified this as the mechanism responsible for upregulating ATR. Hence, ATM and ATR operate both upstream and downstream of each other in their kinase cascades [124, 125].

Matsuoka et al. (2007) identified 700 potential substrates of ATM and ATR by exposing cells to radiation and measuring changes in phosphorylation at ATM/ ATR binding motifs [28]. Interpreting the activation data with gene ontology analysis programs PANTHER (S14) and Ingenuity (Ingenuity Systems), they predicted modes of direct protein interaction [28]. Among these interactions, they found that ATM and ATR likely bind to several repair proteins: ATM interacts with RPA and BRCA1, and ATR interacts with BRCA2 [28]. They also initiate kinase cascades that activate each other [28]. Since ATM and ATR recognize similar amino acid motifs and ATM phosphorylates itself, ATR can recognize and activate the autophosphorylation motif on ATM [28]. ATR does not have this motif, but although ATM likely does not phosphorylate ATR, it can phosphorylate mutual substrates, such as TopBP1, that promote ATR activation [28, 29, 30].

Sharing a substrate motif allows ATM and ATR to promote repair pathways that are classically associated with either kinase. Both kinases phosphorylate H2AX to Ser139H2AX/*γ*H2AX, unwinding the lesion from the chromatin structure to make it accessible to regulatory proteins [4, 5]. ATR promotes end-resected DSB repair; likewise, ATM interacts with NER factors, notably XPC [21]. ATR-deficient cells irradiated with UV light are radiosensitive, suggesting other cell cycle check-points, possibly controlled by ATM, activate in response to UV damage [126].

In this model, we consider three different mechanisms of crosstalk between ATM and ATR. Kinase cascades are the first and most direct mechanism: ATR can phosphorylate ATM, and ATM activates TopBP1, one of the cofactors required for ATM activation. Chromatin unwinding is another: both kinases promote histone modifications that unwind and stabilize damaged sections of DNA, so the involvement of one kinase makes the damaged region more accessible to the other. And lastly, cellular processing can change the structure of a lesion: DSB repair often involves creating RPA-bound ssDNA, and RPA-bound ssDNA can break to form DSBs. By identifying the lesions *γ* radiation and UV damage cause, how these lesions are processed by the cell, and how they interact with ATR and ATM, we simultaneously model these three forms of crosstalk.

## 4 Model

We introduce a mathematical model of ATM and ATR response to *γ* and UV radiation, where ATM and ATR are the two dominant regulatory kinases that promote apoptosis in response to these forms of radiation. We estimate the total pro-apoptotic signal received by the cell as the combined activity of ATM and ATR.

Figure 1 outlines structural changes of the lesions represented in the model. The first column in Figure 1 shows the four types of DNA lesion we consider in this model: simple DSBs, complex DSBs, multiply damaged sites, and UV photoproducts. Additional classes in Figure 1 show changes in lesion structure as they undergo repair or induce replication stress, naming the repair process and key proteins responsible for each change.

**Figure 1:**
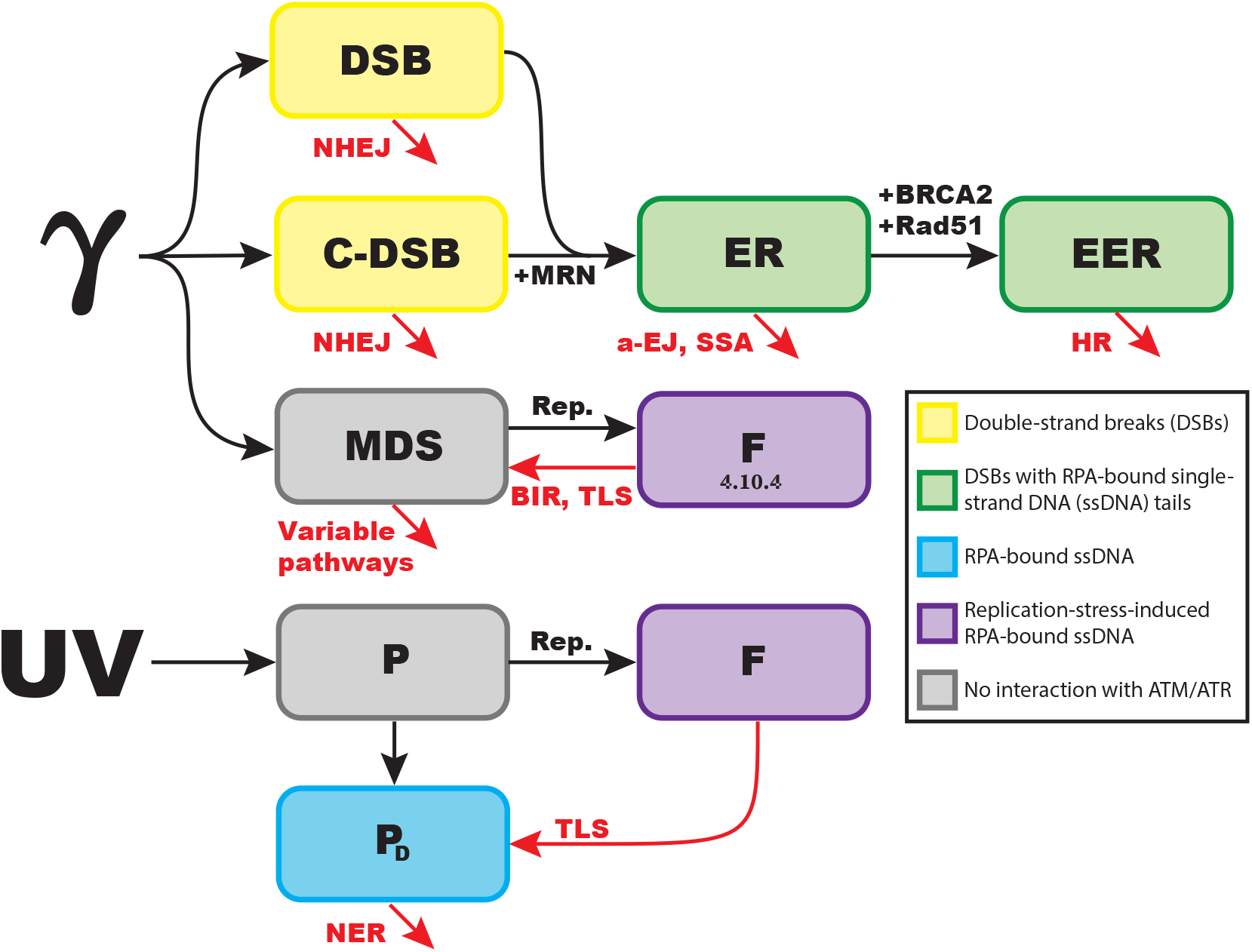
The possible fate of each lesion class in the model. **GAMMA RADIATION:** Gamma radiation creates simple and complex double-strand breaks (DSBs, C-DSBs) and clusters of damaged bases (multiply damaged sites, MDS). Non-homologous end joining (NHEJ) can repair DSBs. If end-resection enzymes outcompete NHEJ factors, DSBs undergo end resection and move to the end-resected DSB class (ER). DSBs in this class can be repaired by alternate end joining (a-EJ) or single-strand annealing (SSA). Regulatory proteins such as BRCA2 and Rad51 may induce further end-resection, which moves the DSB into the extensive-end-resected class (EER) where it can be repaired by homologous recombination (HR). **UV RADIATION:** UV radiation creates photo-products, which may appear on either transcribed (P_*T*_) or non-transcribed (P_*N*_) DNA. Once a photoproduct is detected (*P_D_*), it undergoes nucleotide excision repair (NER). **REPLICATION STRESS:** If the replication machinery encounters an existing lesion, it creates a stalled fork *F*. A stalled fork may be repaired by translesion synthesis (TLS), break-induced repair (BIR), or homologous recombination (HR). These processes use TLS polymerases to pair bulky lesions with complementary bases, leaving the original lesion intact. The 4.10.4 label is used in a later section. Black arrows denote lesion creation, progression, or detection; red arrows represent lesion repair.

The processes in Figure 1 are lesion-specific, but interactions between lesions and regulatory proteins, including ATM and ATR, all follow the same generic structure. Section 4.1 contains a generic system of equations that captures this structure. In the following sections, 4.3 and 4.4, we introduce simplifications, structural changes, and repair processes for each type of lesion. Sections 4.5 and 4.6 concern classes of active ATM and ATR, respectively. Section 4.7 covers cell cycle progression and Section 4.8 shows the full ODE system. We then document the process of calibrating the model to 25 qualitative experimental claims (Section 4.9) and show how the model is altered to represent seven simulated experiments (Section 4.10).

### 4.1 Generic lesion class behavior

Once induced, a lesion can change state in five ways: it can bind to ATM, bind to ATR, change lesion classes through processing (e.g., through end resection), undergo repair, or, if the cell is in S phase, stall a replication fork. Because every lesion in the model evolves in the same basic manner, we outline these structural patterns in a system that represents generic lesion behavior. Let *L* be the number of unbound generic lesions. The initial condition 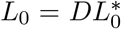 represents the number of lesions *L* induced by *D* Gy of *γ* radiation, where 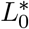 is the estimated number of lesions in class *L* induced by 1 Gy of *γ* radiation. For clarity, we color-code the terms in these equations according to which of the five processes they represent (Table 1).

**Table 1:** Color-coding for state variable equations.

| Purpose of term |
| --- |
| ATM binding |
| ATR binding |
| Lesion state transitions |
| Repair (for lesions) or dephosphorylation (for ATM and ATR) |
| Replication stress |

**Table 2:**
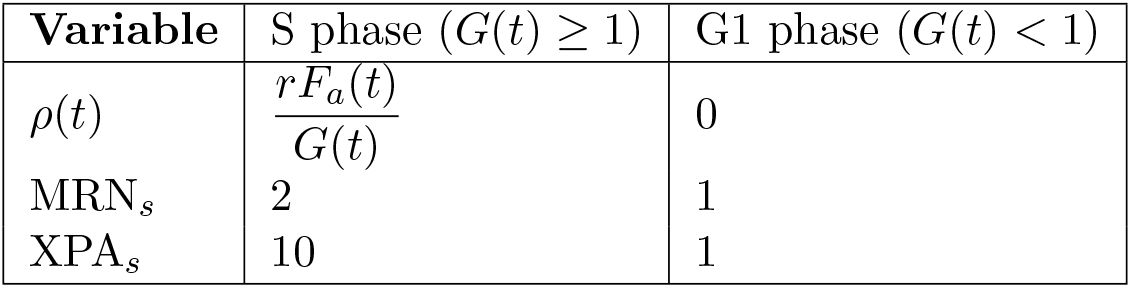
Cell-cycle-specific parameters.

Figure 2 graphically relates the components of the generic lesion system, including active ATM and ATR classes.

**Figure 2:**
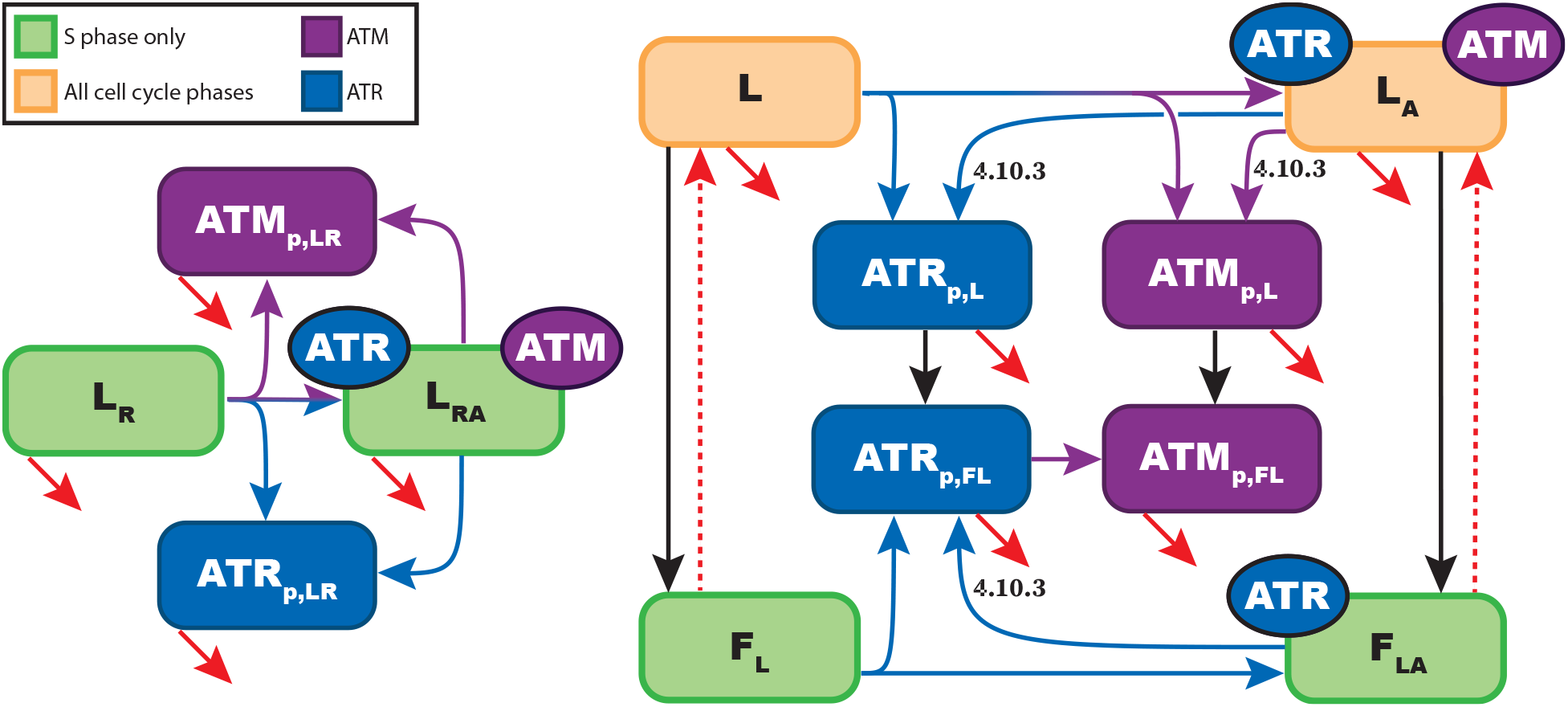
Diagram of state variable changes in the generic lesion system. Red arrows represent repair processes; black arrows represent replication fork stress. The red dashed arrows are optional interactions for when replication fork repair leaves the original lesion intact. Lesion classes with green backgrounds only change during S phase, and lesion classes with orange backgrounds change during all phases of the cell cycle. Purple and blue arrows represent ATM and ATR binding, respectively, and ATM and ATR classes are color-coded in the same manner. A split blue-and-purple arrow indicates that a lesion can interact with either ATR or ATM, but not necessarily both, to enter the kinase-bound class. The 4.10.3 label refers to a later section.

In the equations below, the function 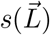 represents transitions between *L* and all other lesion classes, where 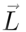 is the set of all lesion classes. Generic lesions undergo repair at a constant rate *k_r_*. Lesions bound to active ATM or ATR undergo modifications, such as histone unwinding, that make them more accessible to regulatory proteins (Section 3.4.3). We separate lesions that have interacted with at least one ATM or ATR molecule into a kinase-bound class (*L_A_*). For kinase-bound lesions, we assume ATM or ATR induces chromatin unwinding on a faster timescale than DNA repair. Because unwinding makes these lesions more accessible to nucleic proteins than unbound lesions, we assume all protein interactions—including end resection, repair, and, notably, ATR/ATM recruitment—happen faster. We represent chromatin restructuring by a constant scaling factor, *r_A_ >* 1, such that regulatory proteins interact with kinase-bound lesions *r_A_* times faster than they do with unbound lesions. We assume this rate is independent of the number of kinases bound to the lesion: as long as the lesion has interacted with at least one kinase, it is permanently altered, even if all kinases dissociate afterward. Two functions, 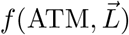 and 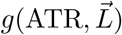, govern ATM and ATR binding, respectively.

While *L_A_* and *L_AR_* compartmentalize lesions that have interacted with ATM or ATR at least once, ATM_*p,L*_ and ATR_*p,L*_, the phosphorylated kinase classes, represent the total number of bound kinases across all lesions of type *L*. Kinases deactivate either spontaneously, at a basal dephosphorylation rate *k_dAT R_* or *k_dAT M_*, or when the lesion they are bound to undergoes repair.

During S phase, we assume replication machinery encounters lesions at an average rate *ρ*. If we induce damage in the middle of S phase, only lesions on unreplicated DNA can cause replication stress. We therefore introduce two additional kinase-unbound and -bound lesion classes, *L_R_* and *L_AR_*, with associated kinase classes ATM_*p,LR*_ and ATR_*p,LR*_. These lesions do not cause replication stress but otherwise behave identically to their unreplicated counterparts. When the cell is not in S phase, *ρ* = 0. Figure 2 separates the evolution of unreplicated lesion classes, which also represent lesion behavior outside of S phase, from the replication stress subsystem.

Replication forks stall when they separate a lesion from its complementary strand. The RPA-bound ssDNA in these stalled forks directly interacts with ATR, which may then activate ATM. Stalled forks undergo repair at rate *k_L_* and break endogenously, forming DSBs, at rate *k_P L_*. The process that repairs the stalled fork may not repair the original lesion; if this is the case, it returns the lesion to its original class. Fork-associated ATR behaves the same way it does when bound to *L*; fork-associated ATM binds to ATR at a rate *k_aa_f* (ATM). Even if the original lesion is still present, all ATM and ATR proteins dissociate from the stalled fork upon repair. We let *F_L_* be the number of replication forks stalled at lesions of type *L*, *F_LA_* be the number of ATR-bound replication forks of this type, and 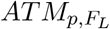 and 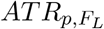 be the number of ATM or ATR proteins bound to these forks.

Lastly, let ATM_*tot*_ be the total number of ATM molecules in the cell, which we assume is constant, and let ATM_*p*_ be the number of phosphorylated ATM molecules at time *t*. The difference between these quantities is the number of free ATM molecules at time *t*. Free ATR is the difference between analogous quantities ATR_*tot*_ and ATR_*p*_. The system that governs the kinase-bound classes, replication stress classes, and associated kinase classes of a generic lesion is as follows:

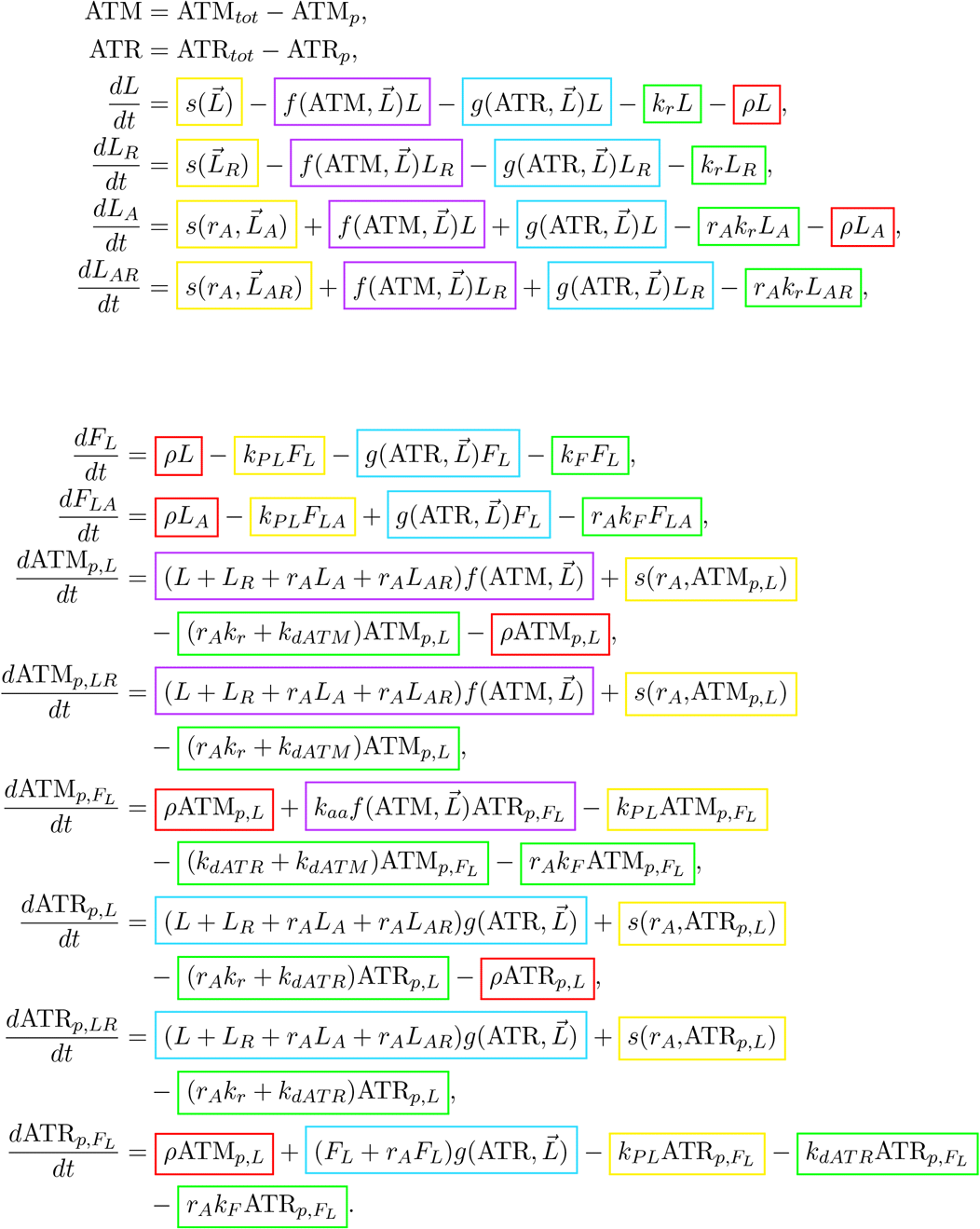

While a model built according to these rules would need twelve state variables for each type of lesion, some classes can be removed without significantly changing the model’s behavior. The following sections outline differences between the generic class and specific lesions, as well as introducing state changes between lesion types.

### 4.2 Initial conditions

The state variables used in this system can be split into five categories: active ATM, active ATR, lesions that have been processed by the cell, lesions that have yet to be processed, and cell cycle control variables. We expose the cell to radiation at *t* = 0. In an undamaged cell, all variables in the first three categories start at zero.

To estimate the number of lesions produced by radiation exposure, we use measurements from the existing literature. Meldrum et al. (2003) state that 4500 photoproducts are produced per J/m^2^ of UV radiation the cell receives [59]. For *γ* radiation, we use Ward’s 1988 meta-study, which finds that each Gy of *γ* radiation produces four complex DSBs, 36 simple DSBs, and 400 MDS per cell [45]. Both papers model the dose-response relationship as linear for lower levels of damage, noting that both forms of radiation reach a threshold above which complex combinations of lesions start to appear at a higher frequency. Our model is not capable of simulating lesion interactions beyond this threshold; its results should only be considered relevant for radiation exposure under 50 J/m^2^ UV or 100 Gy of *γ* radiation.

Lesions induced during S phase can cause replication stress only if the DNA they appear on has not already been replicated. Let *s* represent the proportion of the genome that has been replicated by *t* = 0 and let *G_tot_* refer to the total length of the genome, approximately 3.3 × 10^9^ bp in humans [127]. If *L* lesions are uniformly distributed throughout the genome, we expect radiation to induce approximately (1 − *s*)*L* lesions on unreplicated DNA.

Radiation given to a cell during S phase can damage not only the cell’s original genome but the newly replicated DNA as well. Hence, the cell cycle may influence how many lesions the cell accumulates after exposure to a damaging agent. Radiation damages DNA by transferring free energy to base pairs, a process we expect to occur at a rate independent of DNA density. A cell irradiated during S phase would then have more total lesions than an irradiated non-replicating cell, but the lesion density would be the same. An actively replicating cell has roughly (1 + *s*)*G_tot_* base pairs, and if the lesion density estimates above originate from dormant cells, we expect radiation to induce *L*(1 + *s*) lesions. Contrast this with chemical damaging agents, which have a limited reaction rate: if these agents catalyze DNA at close to their maximal speed in a non-replicating cell, we would expect an actively replicating cell to have the same number of lesions at a lower density.

Lastly, we estimate the total number of lesions by multiplying the above lesion densities by the magnitude of induced damage, *D_IR_* Gy for *γ* radiation and *D_UV_* J/m^2^ for UV. Under these assumptions, the initial number of lesions in each compartment is

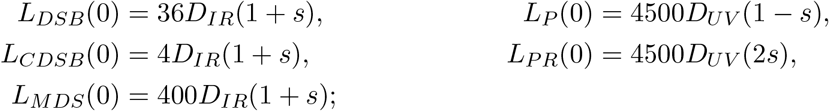

where *G*(0) = *G_tot_*(1 − *s*). We set all other initial quantities to 0, except for Pol*δ_f_*, which we set to its steady state in an undamaged, non-replicating cell.

### 4.3 γ-radiation-induced lesions

This model includes four classes of double-strand break: simple DSBs (*L_DSB_*), complex DSBs (*L_CDSB_*), MRN-processed end-resected DSBs (*L_ER_*), and extensively end-resected Rad51-bound DSBs (*L_ERR_*), in addition to one class for multiply damaged sites (MDS). Figure 3 shows how these lesion classes evolve.

**Figure 3:**
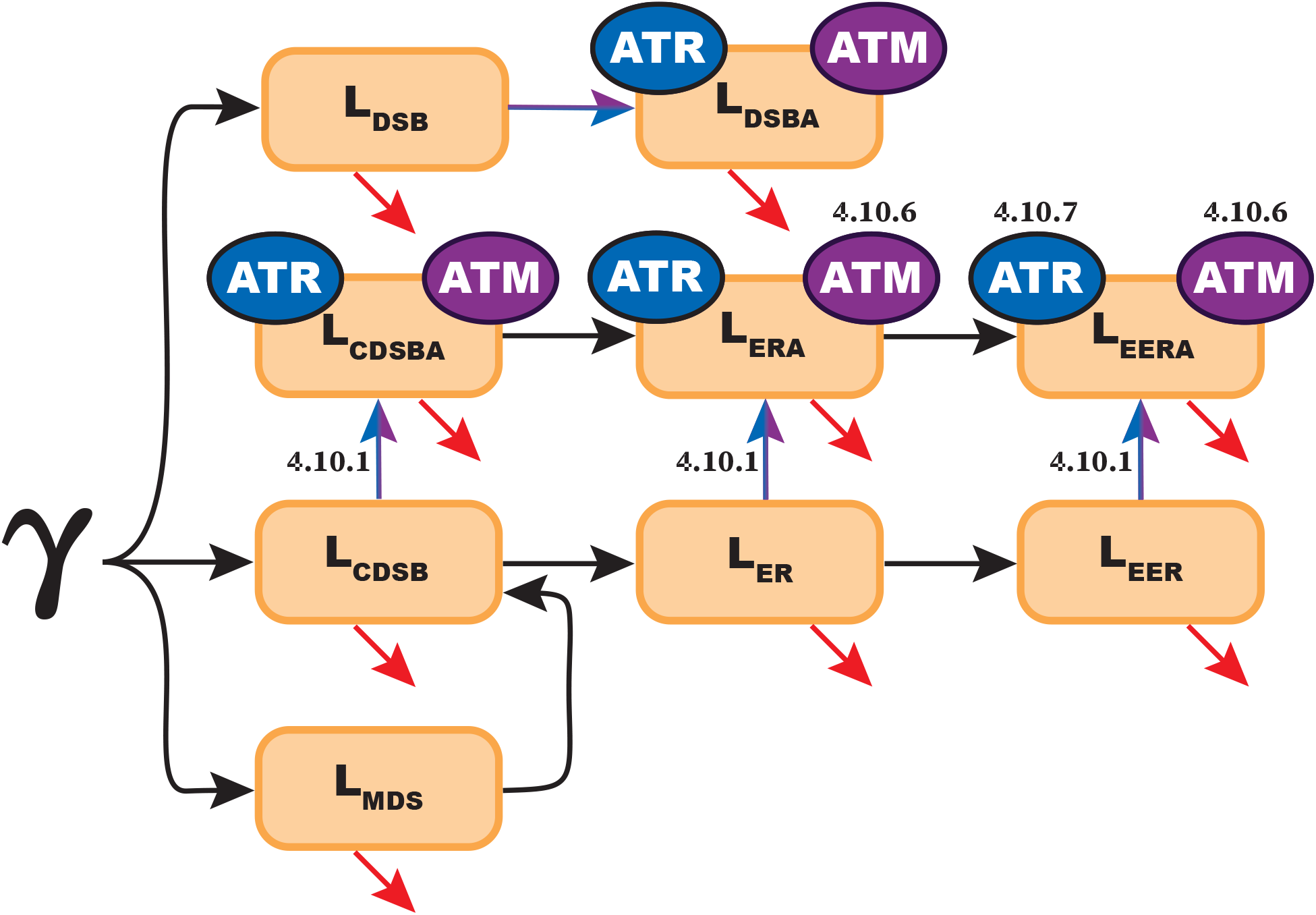
The evolution of *γ*-radiation-induced lesion classes. Gamma radiation induces simple DSBs (DSBs), complex DSBs (CDSBs), and multiply damaged sites (MDSs). Once created, these lesions may bind to ATM or ATR, be repaired, or be processed by the cell to create secondary lesions: end-resected DSBs (ER) or extensively end-resected DSBs (ERR). Arrows in purple or blue denote interactions with ATM or ATR, respectively. The 4.10 labels are used in a later section.

Unprocessed DSBs, both simple (*L_DSB_*) and complex (*L_CDSB_*), behave similarly to the generic lesion class. They are canonically associated with ATM but can bind to either ATM or ATR before end resection (Section 3.4.1). NHEJ repairs unprocessed DSBs, where complex DSBs take longer to repair (*k_NHEJ_ > k_SNHEJ_*, Section 3.3.3). Unprocessed DSBs move to a different lesion class by being resected by MRN complex, and endemic ssDNA breakage can convert MDS to DSBs. *L_DSB_* and *L_CDSB_*, along with their kinase-bound counterparts *L_DSBA_* and *L_CDSBA_*, evolve as follows:

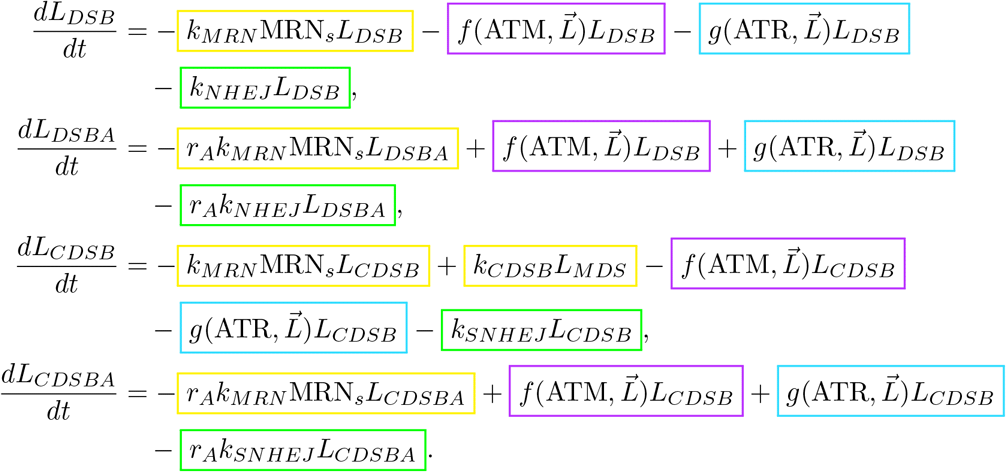

MRN_*s*_ represents the factor by which MRN binding rate increases during S phase. MRN_*s*_ can be set to 0 to model an MRN-knockout cell. We define the two unspecified functions that control ATM and ATR activation, 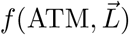 and 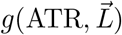, in the sections that describe ATM and ATR activation (Sections 4.5 and 4.6).

DSBs in either class that have been resected by MRN complex move to the end-resected class, *L_ER_*. These end-resected DSBs are repaired at a low rate by SSA or undergo BRCA2-mediated Rad51 strand invasion and move to the extensively end-resected class, *L_EER_*, representing DSBs that have undergone the extensive end resection required for HR repair (Section 3.3.3). Longer regions of ssDNA break at a low rate (*k_P L_*) to form DSBs with long ssDNA tails. We assume DSBs in the extensively end-resected class are still capable of activating ATR (Section 3.4.1). The end-resected DSB classes, including kinase-bound classes *L_ERA_* and *L_EERA_*, evolve as

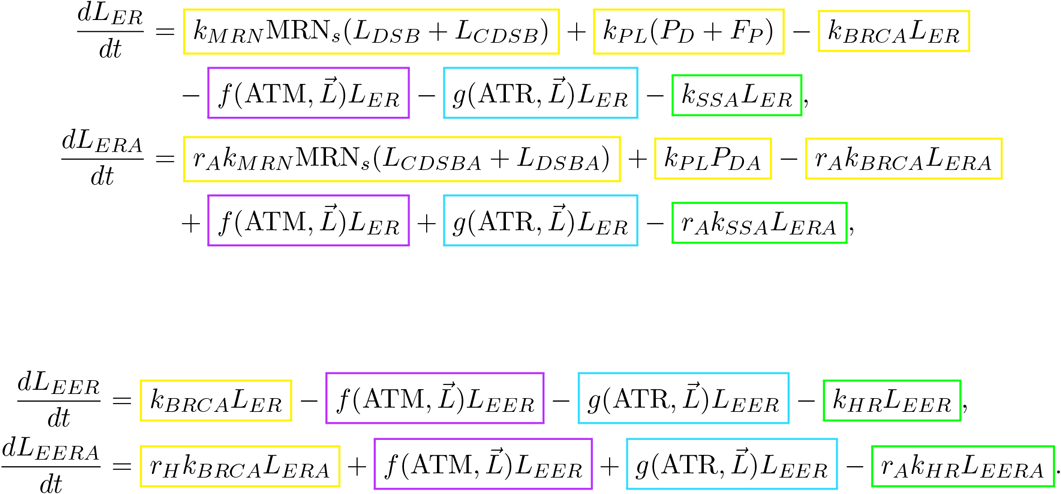

Gamma radiation also creates multiply damaged sites (MDS). Without knowing how the structure of these lesions influences repair, we cannot accurately model ATR involvement in the repair of MDS (Section 3.3.2). We therefore overlook the possibility that MDS interacts directly with ATR or ATM and omit the kinase-bound MDS classes. The two kinases instead interact with DSBs created as a product of improper MDS processing. The MDS class, *L_MDS_*, is directly solvable:

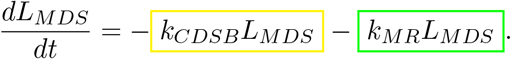

If the cell is exposed to *γ* radiation during replication, *γ*-radiation-induced lesions may cause replication stress. We alter the model to account for this possibility in Section 4.10.4.

### 4.4 UV photoproducts

Photoproducts induce RPA-bound ssDNA as either a repair intermediate or a result of replication stress (Section 3.3.4). We separate undetected photoproducts into an unreplicated class (*L_P_*) and a replicated class (*L_P R_*), where undetected photoproducts in the unreplicated class can stall replication forks:

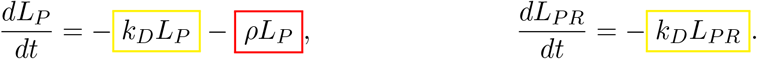

The equation governing *L_P R_* is solvable by hand. There was no noticeable difference in dynamics between this model and an alternate version that accounts for the difference in speed between transcription-coupled and global-genome NER (Section 3.3.4, results not shown).

Once the cell detects photoproducts in any of these four initial classes, they move into the detected class, *L_P D_*. The cell repairs detected photoproducts by NER, which creates RPA-bound ssDNA as an intermediate. Free ATR can bind to this ssDNA (Section 3.4.2). If this happens before the photoproduct is repaired, the repair intermediate moves to the ATR-bound class, *L_P DA_*:

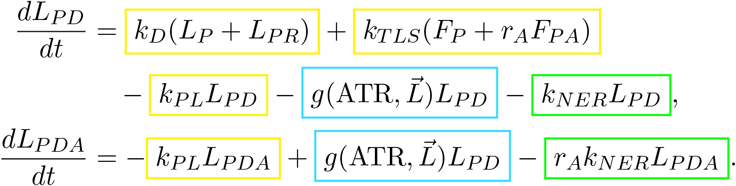

We define the two unspecified functions that control ATM and ATR activation, 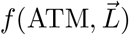 and 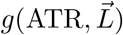, in the sections that describe ATM and ATR activation (Sections 4.5 and 4.6).

We represent forks stalled on photoproducts by *F_P_* and *F_P A_*. Recall that TLS, the process that synthesizes complementary DNA at stalled replication forks, does not repair photoproducts. Once TLS is complete, the remaining photoproduct moves into the *P_D_* class. There is no class for replication forks stalled on detected photoproducts: the NER excision site is flanked by helper proteins that block replication machinery. *F_s,P_* and *F_s,P A_* evolve according to

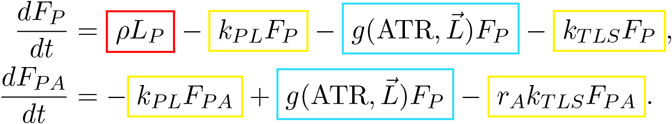

Any ssDNA in the above classes has a low chance of breaking to become a DSB. When this occurs, the resulting lesion moves to the end-resected class that preserves its kinase-bound status, either *L_ER_* or *L_ERA_*. Figure 4 shows a diagram of this system.

**Figure 4:**
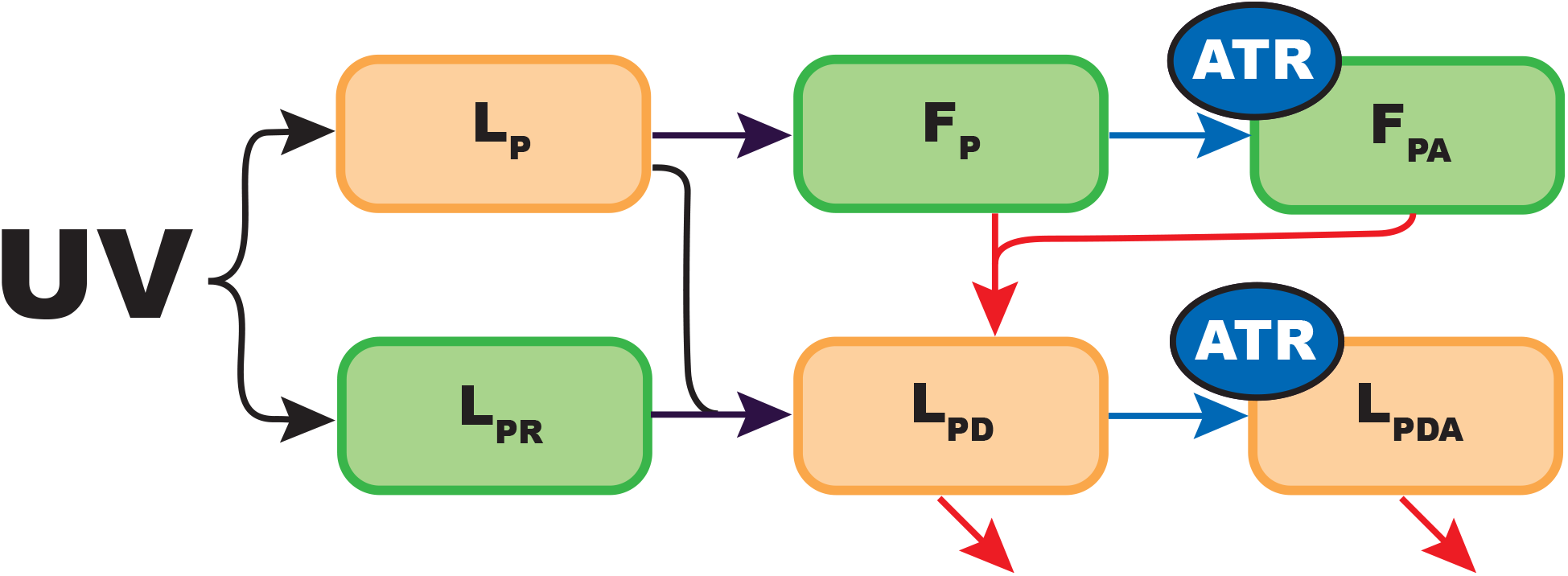
Diagram showing the evolution of UV photoproducts. Here, *L_P_* represents undetected photoproducts, *L_P D_* represents detected photoproducts, and *F_P_* represents forks stalled on photo-products. Arrows in purple or blue denote interactions with ATM or ATR, respectively; red arrows represent repair processes. Classes marked in green are only nonzero during S phase.

### 4.5 ATM

ATM is dimeric in its inactive form (Section 3.1). When it encounters a DSB, the two subunits phosphorylate each other and dissociate, with each subunit binding to one side of the break. ATM and other repair pathway proteins then accumulate around the DSB to form a focus (Section 3.4.1). Since hundreds of ATM proteins can bind to a single DSB, the number of active ATM proteins can be larger than the number of DSBs in the cell.

There are four classes of DSB-bound ATM (ATM_*p,DSB*_, ATM_*p,CDSB*_, ATM_*p,ER*_, ATM_*p,EER*_), which follow the same rules of state transfer and repair as their break-class counterparts in Section 4.3. The additional two classes account for ATR-bound ATM at lesions with RPA-bound ssDNA, ATM_*p,P*_ for photoproducts and ATM_*p,F*_ for stalled replication forks (Section 3.4.3). We assume TLS removes all RPA-bound ssDNA from the damaged site, which deactivates all fork-associated ATR and ATM. Overall, the active ATM classes behave as follows:

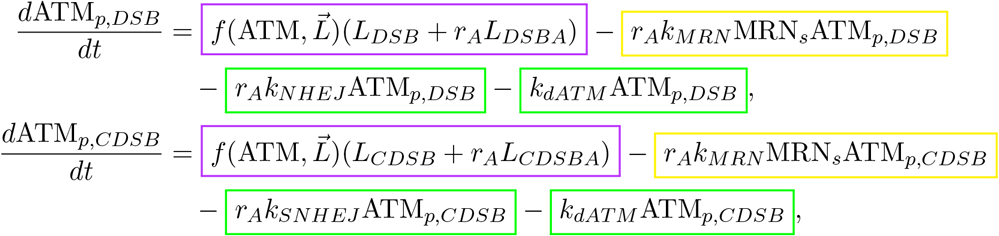

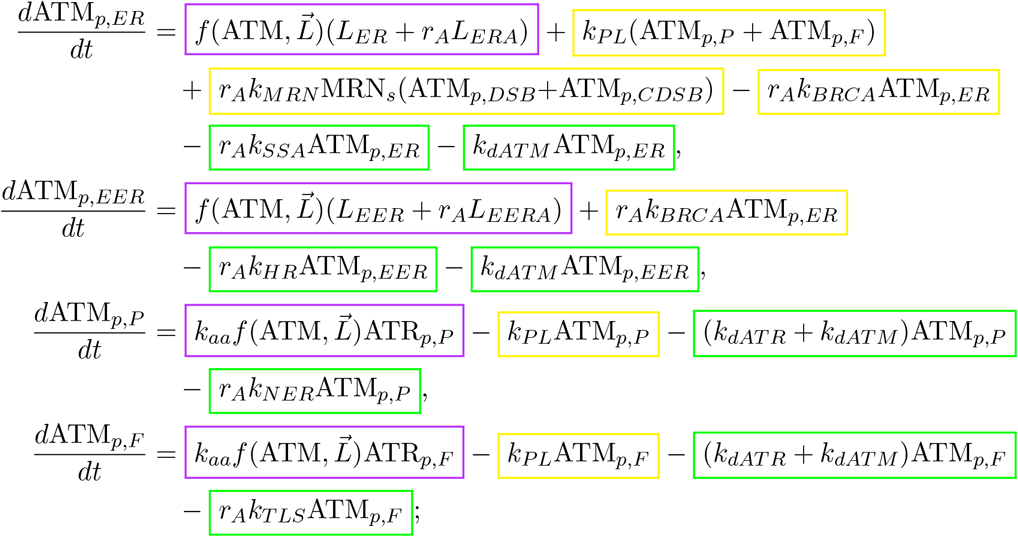

where

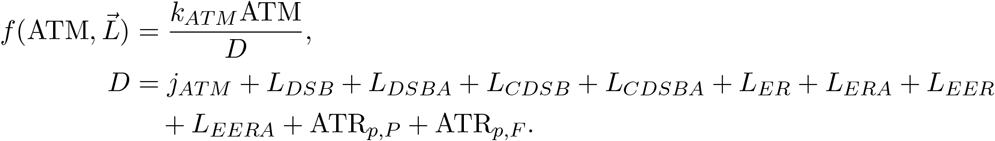

The total number of active ATM proteins per cell is twice the sum of all active classes:

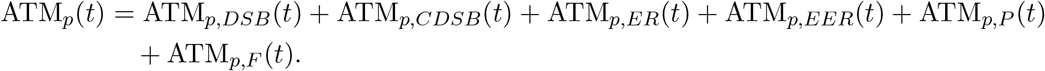

Because ATM splits into two monomers when phosphorylated (Section 3.1), this number can be doubled; here, we omit this scaling factor to avoid confusion. Each rate that affects active ATM, we understand as acting on pairs of active monomers. We assume the total amount of ATM in the cell is held constant on this timescale and calculate the number of free ATM proteins using the conservation equation

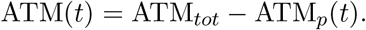

Since we designed this model to act as the upstream module of a larger p53 regulatory system, it does not include p53-dependent ATM regulation. In particular, p53 upregulates transcription of Wip1, which has been shown to suppress ATM activation within 2 hours of damage exposure [128]. With the exclusion of Wip1 and other p53-dependent regulators, ATM behaves in this model as it would in a cell with transcriptionally defective p53. The equations above implicitly assume that ATM quickly re-dimerizes after being dephosphorylated. Figure 5 shows state transitions between ATM and ATR classes.

**Figure 5:**
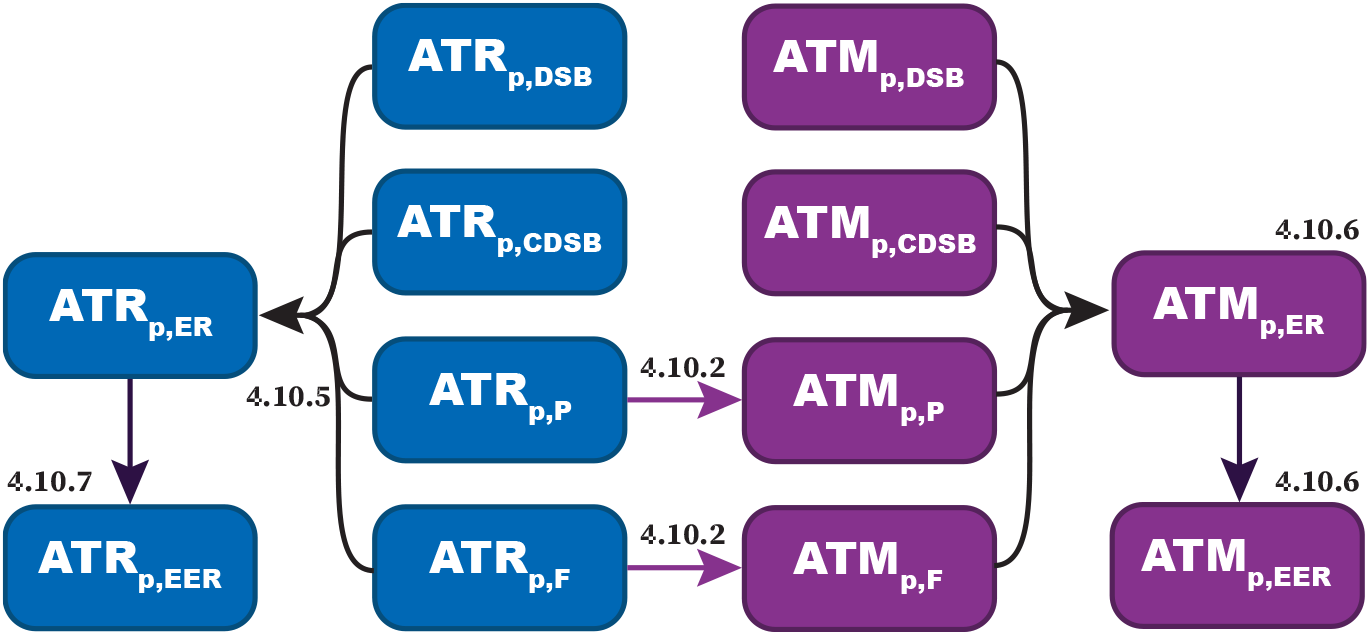
Diagram showing ATM and ATR state changes. With two exceptions, each class of ATM or ATR grows when unbound proteins interact with the lesions indicated in their subscripts. ATM_*p,P*_ and ATM_*p,F*_ instead bind to ATR molecules in their corresponding class. The transitions between kinase classes mirror the transitions between their associated lesions. Arrows in purple denote interactions with ATM. The 4.10 labels are used in a later section.

### 4.6 ATR

RPA-bound ssDNA activates ATR. Our model includes ssDNA created as a result of replication fork stalling (*F_MDS_*, *F_P_*), ssDNA overhangs on DSBs (*L_DSB_*, *L_CDSB_*, *L_ER_*, *L_ERR_*), and ssDNA created as an NER repair intermediate (*L_P D_*). While ATR is associated with stalled replication forks because it is primarily active during S phase, we assume it can be activated by ssDNA from any of these sources. In particular, RPA is involved in NER, which may influence early ATR activation independently of replication stress. We assume RPA binds rapidly to newly created ssDNA; hence, we do not include classes of RPA-unbound ssDNA and allow ssDNA to bind to ATR as soon as it is created [125].

Unlike DSB foci, which have no intrinsic size limit, the length of RPA-coated ssDNA strands limits the number of available ATR binding sites. Our model does not estimate the number of ATR binding sites on each ssDNA strand. Instead, other limiting factors—such as the total amount of ATR and the average time a lesion persists before repair—indirectly constrain the amount of ATR that can bind to a break. Both ATM and ATR activate TopBP1, an essential binding factor between ATR and RPA-bound ssDNA. To account for ATM upregulation of ATR through TopBP1, we identify the terms representing ATR binding to kinase-bound DSBs and multiply them by a factor *k_top_ >* 1.

ATR is most active during S phase and controls S/G2 damage checkpoints. We assume this increase in ATR activity is controlled by cell-cycle-specific regulators, particularly XPA (Section 3.1). Here, we treat ATR cell-cycle-specificity as a piecewise property, where XPA is a constant that scales ATR binding rates and is set to different values for S and G1 phases:

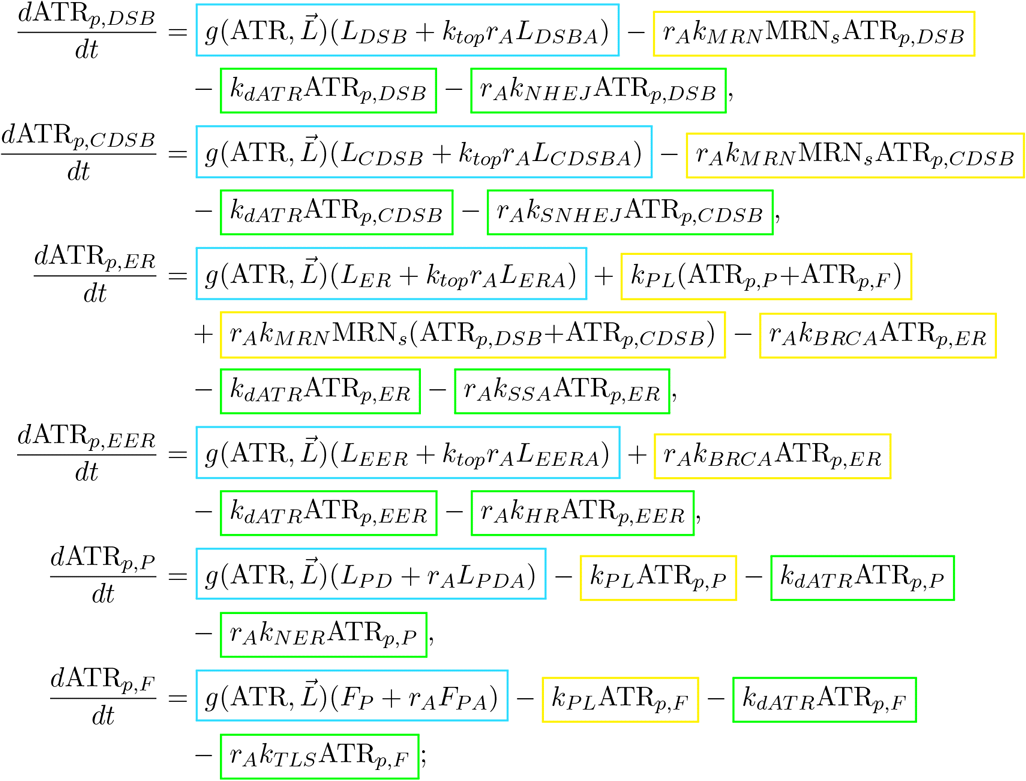

where

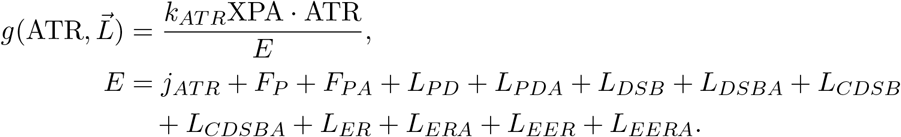

The total number of active ATR proteins in the cell is the sum of all ATR-RPA-ssDNA complex classes, such that

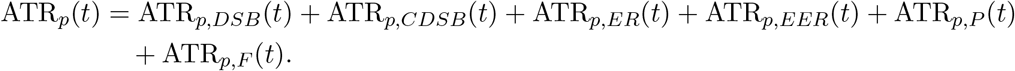

Assuming the total amount of ATR in the cell remains relatively constant, we compute the amount of free, inactive ATR by subtracting the sum of all active ATR classes from the total:

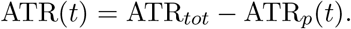

### 4.7 Cell cycle progression

This model supports two parts of the cell cycle: G1 phase and S phase. To model progression through S phase, we record the number of base pairs that have yet to be replicated (*G*) and the amount of free Pol *δ*.

Let *G_tot_* = 3.3 × 10^9^ be the total length of the human genome and let *G*(*t*) be the amount of unreplicated DNA remaining in the cell during S phase. We set *G*(0) = 0 to simulate G1 phase, *G*(0) = *G_max_* to start S phase. We consider S phase to have ended when *G*(*t*) < 1; that is, when less than one base pair remains to be replicated.

We assume origins of replication are distributed uniformly across the genome, so that when *G*(*t*) unreplicated base pairs remain, the ratio between free origins of replication at time *t* and total origins of replication in the genome is approximately 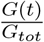. Free Pol *δ* binds with high affinity to these origins of replication during S phase. If ATR is active, it suppresses the rate at which origins of replication fire [39], represented here by decreasing the number of active forks:

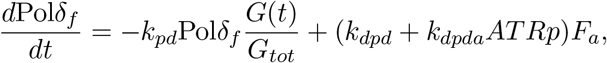

where *F_a_* = Pol*δ_tot_* − Pol*δ_f_* represents the number of active replication forks. The amount of unreplicated DNA, *G*(*t*), can change in two ways: it can be successfully replicated at rate *rF_a_* or become the site of a stalled fork at rate *ρ*:

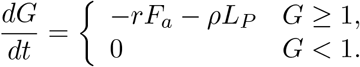

While *r* is held constant, *ρ* varies with lesion density. Here, *r* is the replication fork speed in bp/m, such that *rF_a_* is the net speed of replication by all active forks. If we multiply the number of lesions on unreplicated DNA by *ρ*, the resulting term is the net speed of replication times the lesion density on unreplicated DNA, which estimates the rate at which replication forks encounter lesions. Observe that because both *F_a_* and *G* change over time, *ρ* is not constant. Figure 2 outlines cell-specific function and parameter changes, where the variables assume their S-phase-specific values when *G*(*t*) ≥ 1 and their G1-specific values when *G*(*t*) < 1.

Due to this construction, *G*(*t*) represents the number of available base pairs for replication, not the number of base pairs that must be transcribed before the cell divides. The total amount of unreplicated DNA instead includes *G*(*t*) and all four classes of stalled fork. We model each stalled fork as being 1 bp long.

### 4.8 Full model

We list the evolution of all 29 state variables below.

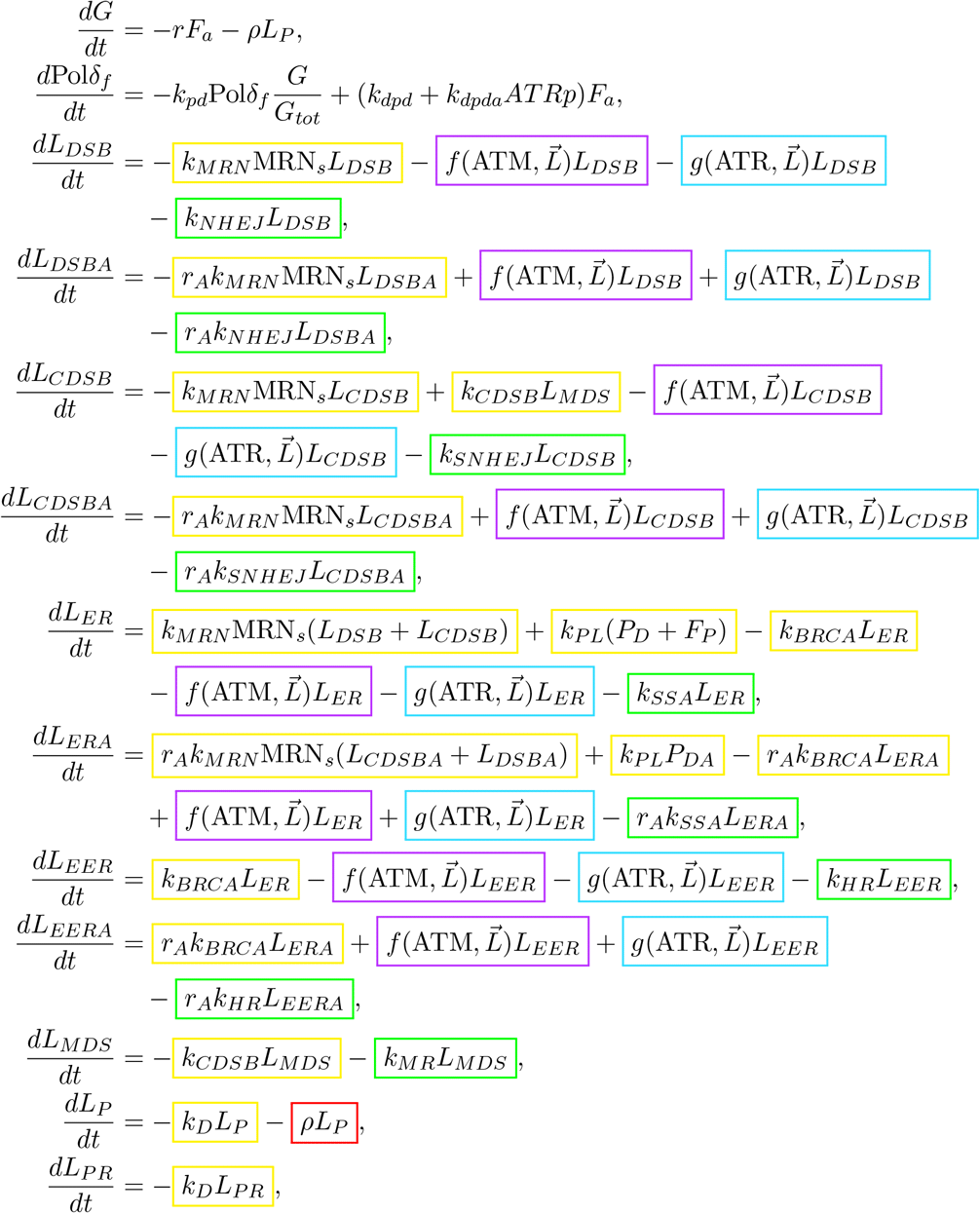

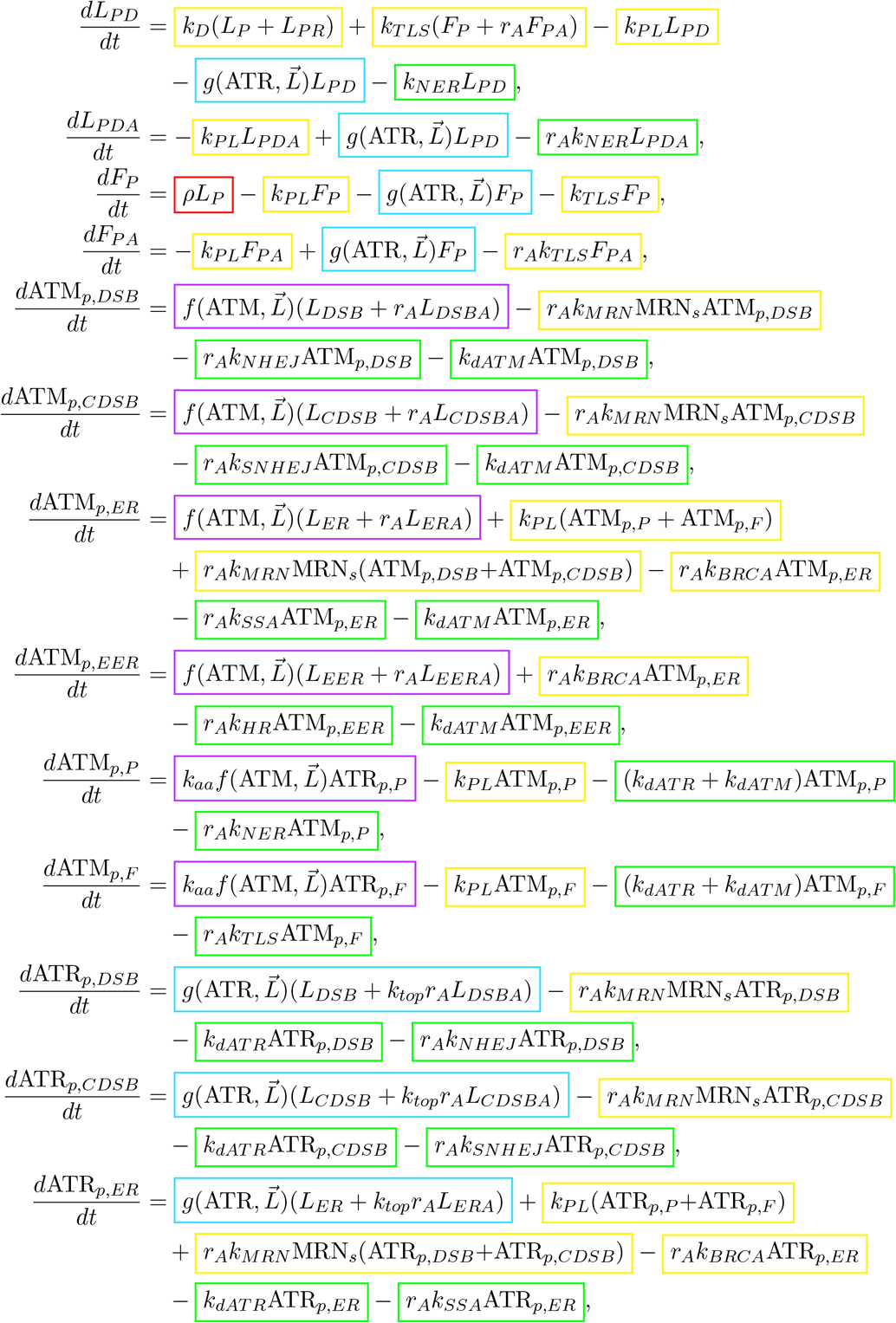

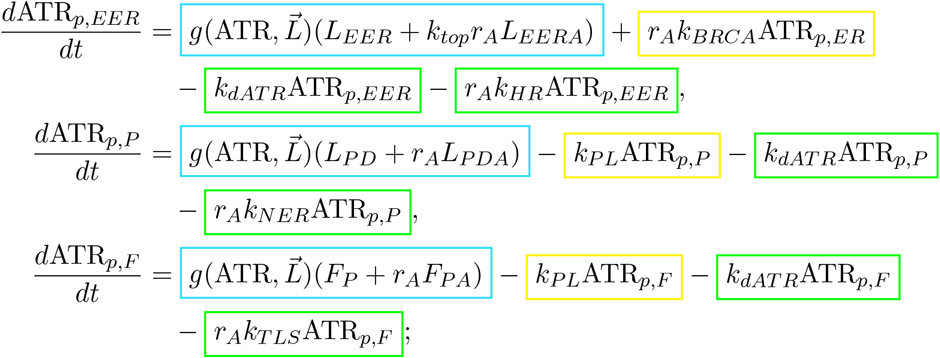

where

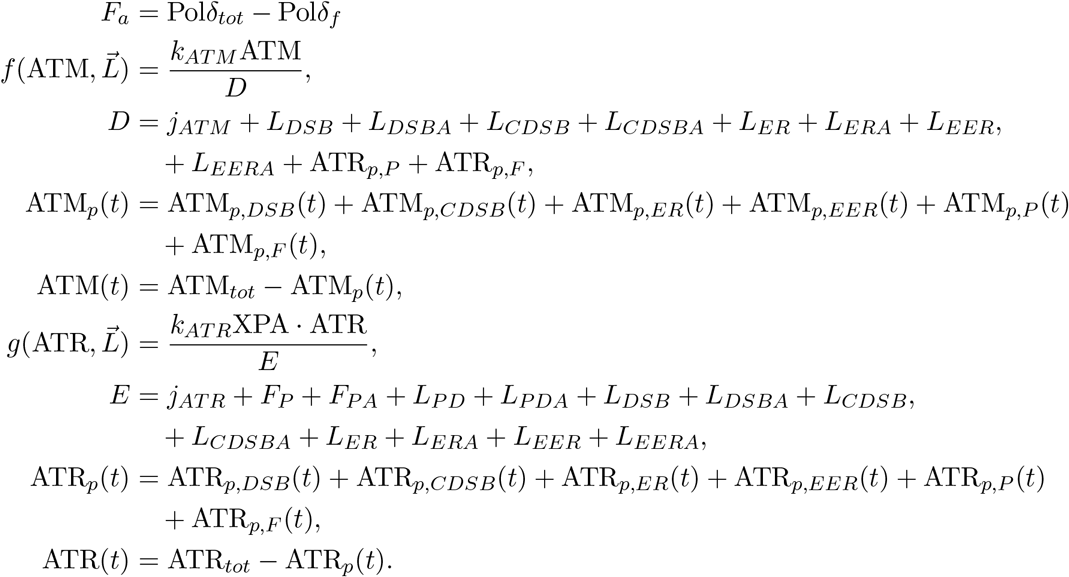

We list the names of each state variable in Table 3.

**Table 3:** State variables used in the ATM/ATR crosstalk model.

| Variable | Description |
| --- | --- |
| $G$ | Number of unreplicated base pairs in genome |
| $\text{Pol}\delta_f$ | Free $\text{Pol}\delta$ available for replication |
| $L_{DSB}$ | Simple double-strand breaks (DSBs) |
| $L_{DSBA}$ | ATM- or ATR-bound simple DSBs |
| $L_{CDSB}$ | Complex DSBs |
| $L_{CDSBA}$ | ATM- or ATR-bound complex DSBs |
| $L_{ER}$ | MRN end-resected DSBs |
| $L_{ERA}$ | ATM- or ATR-bound MRN end-resected DSBs |
| $L_{ERR}$ | Extensively end-resected DSBs |
| $L_{ERRA}$ | ATM- or ATR-bound extensively end-resected DSBs |
| $L_{MDS}$ | Multiply damaged sites (MDS) on unreplicated DNA |
| $L_{MDSR}$ | MDS on replicated DNA |
| $L_{PN}$ | Undetected photoproducts on non-replicated or -transcribed DNA |
| $L_{PNR}$ | Undetected photoproducts on replicated, non-transcribed DNA |
| $L_{PT}$ | Undetected photoproducts on non-replicated, transcribed DNA |
| $L_{PTR}$ | Undetected photoproducts on non-replicated, transcribed DNA |
| $L_{PD}$ | Detected photoproducts |
| $L_{PDA}$ | ATR-bound photoproducts |
| $F_{MDS}$ | MDS-stalled replication forks |
| $F_{MDSA}$ | ATR-bound MDS-stalled replication forks |
| $F_P$ | Photoproduct-stalled replication forks |
| $F_{PA}$ | ATR-bound photoproduct-stalled replication forks |
| $\text{ATM}_{p,DSB}$ | Active ATM bound to simple DSBs |
| $\text{ATM}_{p,CDSB}$ | Active ATM bound to complex DSBs |
| $\text{ATM}_{p,ER}$ | Active ATM bound to MRN end-resected DSBs |
| $\text{ATM}_{p,ERR}$ | Active ATM bound to extensively end-resected DSBs |
| $\text{ATM}_{p,P}$ | Active ATM bound to photoproduct-bound ATR |
| $\text{ATM}_{p,F}$ | Active ATM bound to stalled-replication-fork-bound ATR |
| $\text{ATR}_{p,DSB}$ | Active ATR bound to simple DSBs |
| $\text{ATR}_{p,CDSB}$ | Active ATR bound to complex DSBs |
| $\text{ATR}_{p,ER}$ | Active ATR bound to MRN end-resected DSBs |
| $\text{ATR}_{p,ERR}$ | Active ATR bound to extensively end-resected DSBs |
| $\text{ATR}_{p,P}$ | Active ATR bound to photoproducts |
| $\text{ATR}_{p,F}$ | Active ATR bound to stalled replication forks |

### 4.9 Calibration and validation

Table 4 lists descriptions and values of all non-cell-specific parameters used in the model.

**Table 4:** Parameters in ATM/ATR crosstalk model. If we have not provided a source, the parameter value represents an order-of-magnitude estimate.

| Parameter | Value | Units | Description | Source |
| --- | --- | --- | --- | --- |
| $r_A$ | 6 | None | Histone unwinding factor | [113] |
| $k_{NHEJ}$ | $\frac{\ln 2}{20r_A}$ | $\text{min}^{-1}$ | NHEJ repair rate | [81] |
| $k_D$ | $2 \times 10^4$ | $\text{prot} \cdot \text{min}^{-1}$ | CPD detection rate | |
| $k_{SNHEJ}$ | $\frac{\ln 2}{300r_A}$ | $\text{min}^{-1}$ | NHEJ with preprocessing | |
| $k_{PL}$ | $7 \times 10^{-8}$ | $\text{min}^{-1}$ | ssDNA breakage rate | |
| $k_{MRN}$ | $\frac{\ln 2}{200r_A}$ | $\text{min}^{-1}$ | MRN complex binding rate | |
| $\text{MRN}_s$ | 2 | None | MRN upregulation in S phase | |
| $k_{BRCA}$ | $\frac{\ln 2}{60r_A}$ | $\text{min}^{-1}$ | Rad51 binding rate | |
| $k_{SSA}$ | $\frac{\ln 2}{200r_A}$ | $\text{min}^{-1}$ | Rate of SSA | |
| $k_{HR}$ | $\frac{\ln 2}{600r_A}$ | $\text{min}^{-1}$ | HR rate | [81] |
| $k_{NER}$ | $\frac{\ln 2}{20r_A}$ | $\text{min}^{-1}$ | NER rate | [21] |
| $k_{ATM}$ | 0.5 | $\text{prot} \cdot \text{min}^{-1}$ | Maximal ATM activation rate | |
| $j_{ATM}$ | 4000 | prot | ATM half-maximal activation parameter | |
| $\text{XPA}_s$ | 10 | None | ATR upregulation in S phase | |
| $k_{ATR}$ | 0.03 | $\text{prot} \cdot \text{min}^{-1}$ | Maximal ATR activation rate | |
| $j_{ATR}$ | $4 \times 10^4$ | prot | ATR half-maximal activation parameter | |
| $k_{dATM}$ | 0.005 | $\text{min}^{-1}$ | ATM basal deactivation rate | |
| $k_{dATR}$ | 0.005 | $\text{min}^{-1}$ | ATR basal deactivation rate | |
| $r$ | 3000 | bp/m | Maximal DNA replication rate | |
| $k_{pd}$ | 0.1 | $\text{min}^{-1}$ | Polδ binding rate to DNA | |
| $k_{dpd}$ | 0.005 | $\text{min}^{-1}$ | Basal Polδ dissociation rate | |
| $k_{dpda}$ | $1 \times 10^{-5}$ | $(\text{prot} \cdot \text{min})^{-1}$ | Polδ inhibition by ATR | |
| $k_{TLS}$ | $\frac{\ln 2}{100r_A}$ | $\text{min}^{-1}$ | TLS repair rate | |
| $\text{Pol}\delta_{tot}$ | $3.3 \times 10^3$ | prot | Total Polδ proteins in cell | |
| $k_{aa}$ | 0.008 | None | Rate of ATM activation by ATR | |
| $j_{aa}$ | 4000 | prot | ATM activation by ATR half-maximal parameter | |
| $k_{top}$ | 4 | None | ATR binding to ATM-bound DSBs | |
| $\text{ATM}_{tot}$ | 5000 | prot | Total ATM proteins in cell | [129, 130] |
| $\text{ATR}_{tot}$ | 1000 | prot | Total ATR proteins in cell | [129, 130] |

#### 4.9.1 Cell-specific parameters

We treat total amounts of ATM, ATR, and XPA as constant, cell-type-specific parameters. Ho et al. analyzed the yeast proteasome by converting mass spectrometry measurements of protein abundance to estimates for molecules per cell [129]. They measure 2000–3500 ATM and ATR proteins per cell, with a maximum of 10^4^ and a minimum of 10 molecules per cell.

This provides an order-of-magnitude estimate that may not apply to human cells. In 2019, Wang et al. performed a comparative proteomics study across 29 types of human tissue (Figure 6), which they report in units of m/z, representing mass-to-charge ratio [130]. Their results suggest that ATM is more abundant than ATR in most tissues, sometimes by two orders of magnitude. The exceptions are fat cells and testes, in which they measured roughly equal amounts of ATM and ATR; and liver cells, the only type of cell in which ATR was expressed in greater abundance than ATM. In two types of tissue—the esophagus and heart—Wang et al. detected no expression of ATR.

**Figure 6:**
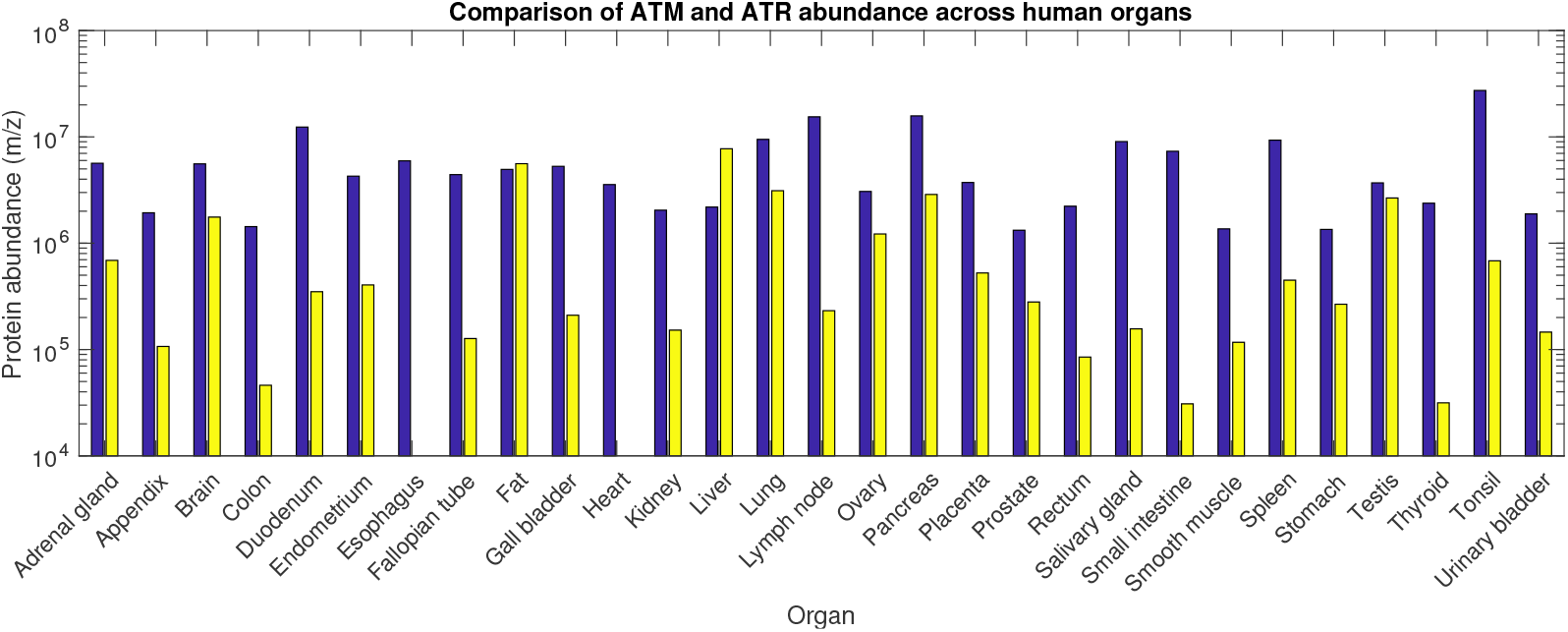
Estimates of protein abundance for ATM and ATR adapted from Wang 2019. Here, blue bars represent ATM abundance in each type of tissue and yellow bars represent ATR.

#### 4.9.2 Damage repair rates

We compared the model to data sets measuring the effects of ATM or ATR silencing on damage repair (Figure 7).

**Figure 7:**
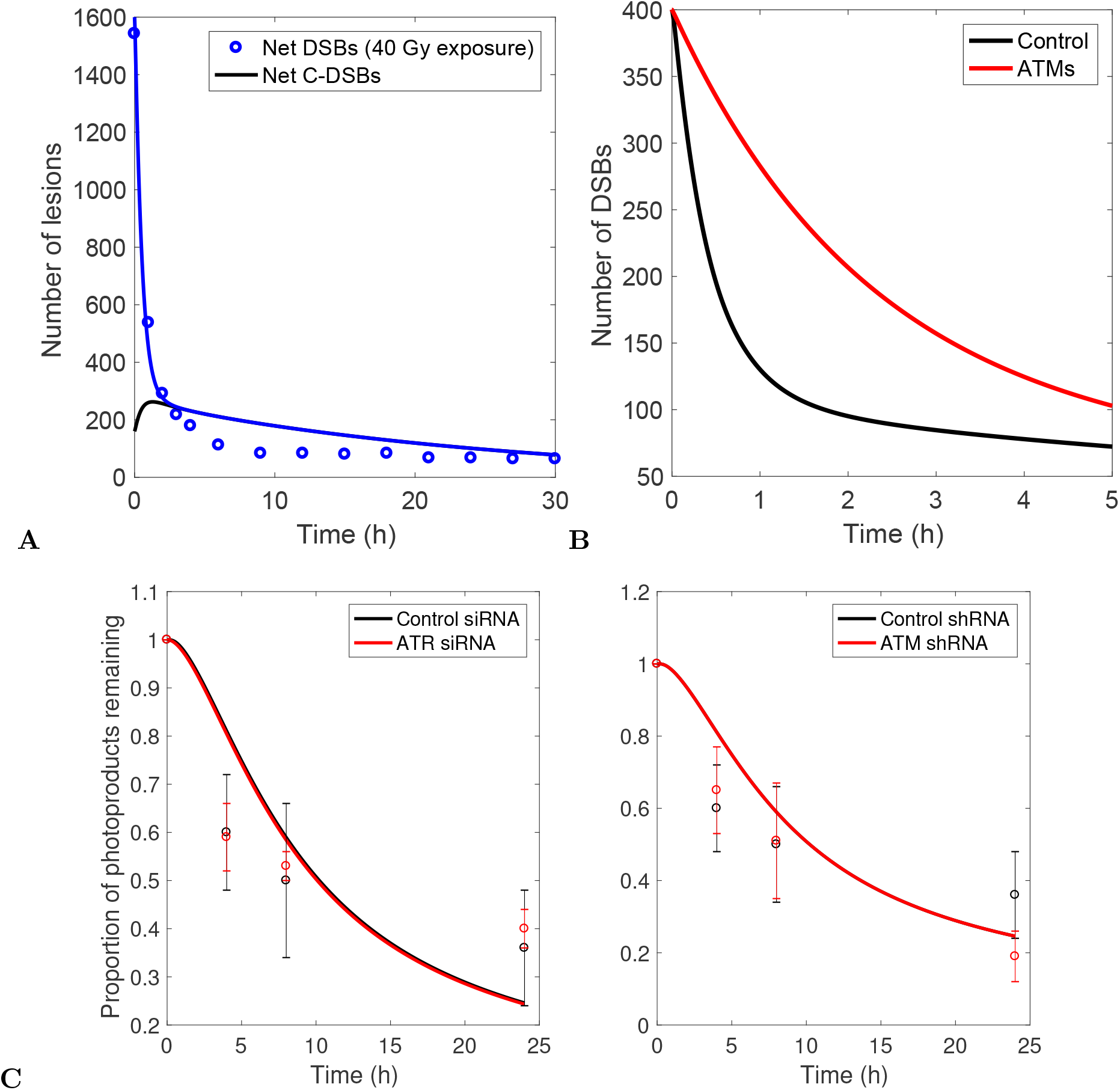
Experiments and data sets used for calibrating damage repair. **(A)** Double-strand break repair kinetics measured by DiBiase et al, 2000 [81]. DiBiase et al. identify a bimodal pattern of DSB repair where the cell’s long-term response to DNA damage depends on a few persistent lesions. **(B)** DSB repair kinetics with and without ATM silencing [113]. Reynolds et al. show that ATM silencing slows DSB repair by a factor of 3.5. **(C**) CPD repair kinetics with either ATR-silencing RNA (left) or ATM-silencing RNA (right) [21]. Ray et al. predict that silencing either kinase does not significantly affect the CPD repair rate.

**Ray et al. 2013 [21]:** Fig. 6 of Ray et al. 2013 shows the percentage of photoproducts remaining after exposure to 10 J/m^2^ UV in G1-arrested OSU-2 cells treated with control RNA, ATR siRNA, or ATM shRNA. We compare our model to their CPD repair data in Figure 7C. Even with perfect silencing and arrest, knocking down either ATM or ATR has a negligible effect on the rate of CPD repair, in agreement with these authors’ conclusions.

**DiBiase et al. 2000 [81]:** Figure 7A shows a comparison of the model to Fig. 3 of DiBiase et al. 2012, which shows biphasic DSB repair in human glioma cell line MO59-K after exposure to 40 Gy of X-rays. Here, we assume 1 Gy of X-rays and 1 Gy of *γ* rays induce similar types and numbers of lesions. We use these data to calibrate DSB repair rates. DiBiase et al. also estimate the half-life of simple DSBs as 22 minutes and the half-life of complex DSBs as 12 hours.

**Reynolds et al. 2012 [113]:** Fig. 6 of Reynolds et al. shows the relative fluorescence intensity of Ku80, used as a DSB flag, in Ku80-EGFP-tagged Chinese hamster cells exposed to NIR microbeam irradiation. Without knowing the number and type of DNA lesions generated by NIR microbeam irradiation, we cannot qualitatively generate the initial conditions to compare the model to these data. However, the authors estimate that inhibiting ATM extends the average half-life of a DSB from 30 ± 14 minutes to 107 ± 18 minutes, by a factor of 3.5. We achieve this by setting *r_A_* = 6 (Figure 7B).

Table 5 summarizes the claims outlined in this section.

**Table 5:** Comparison of the model results to crosstalk data, part 1.

| Source | Claim | Matched<br>by<br>Model? | Notes |
| --- | --- | --- | --- |
| [21] | Knocking down ATM or ATR does not affect the rate of CPD repair | Yes | Used Fig. 6 for calibration |
| [81] | DSB repair is biphasic | Yes | Used Fig. 3 for calibration |
| [113] | Inhibiting ATM slows complex DSB repair by a factor of 3.5 | Yes |  |

### 4.9.3 ATM/ATR crosstalk

In addition to damage repair, we also calibrated the model to match observations about ATM and ATR activation after exposure to either *γ* or UV radiation.

**Jazayeri et al. 2006 [18]:** This is one of the earliest and most influential papers on ATM/ATR crosstalk. In HeLa and U2OS cells, both ATM and ATR are recruited to *γ*-radiation-induced DSBs within 10 minutes, where ATR works downstream of DSB end resection. Since cells irradiated in G1 did not have activated Chk1, the authors conclude that ATR only responds to DSBs in S and G2 phases. Later works, such as Gamper et al. 2013, report that ATR is weakly activated in G1 phase in response to *γ* radiation [20]. Figure 8A shows the model’s response to a high dose of *γ* radiation.

**Figure 8:**
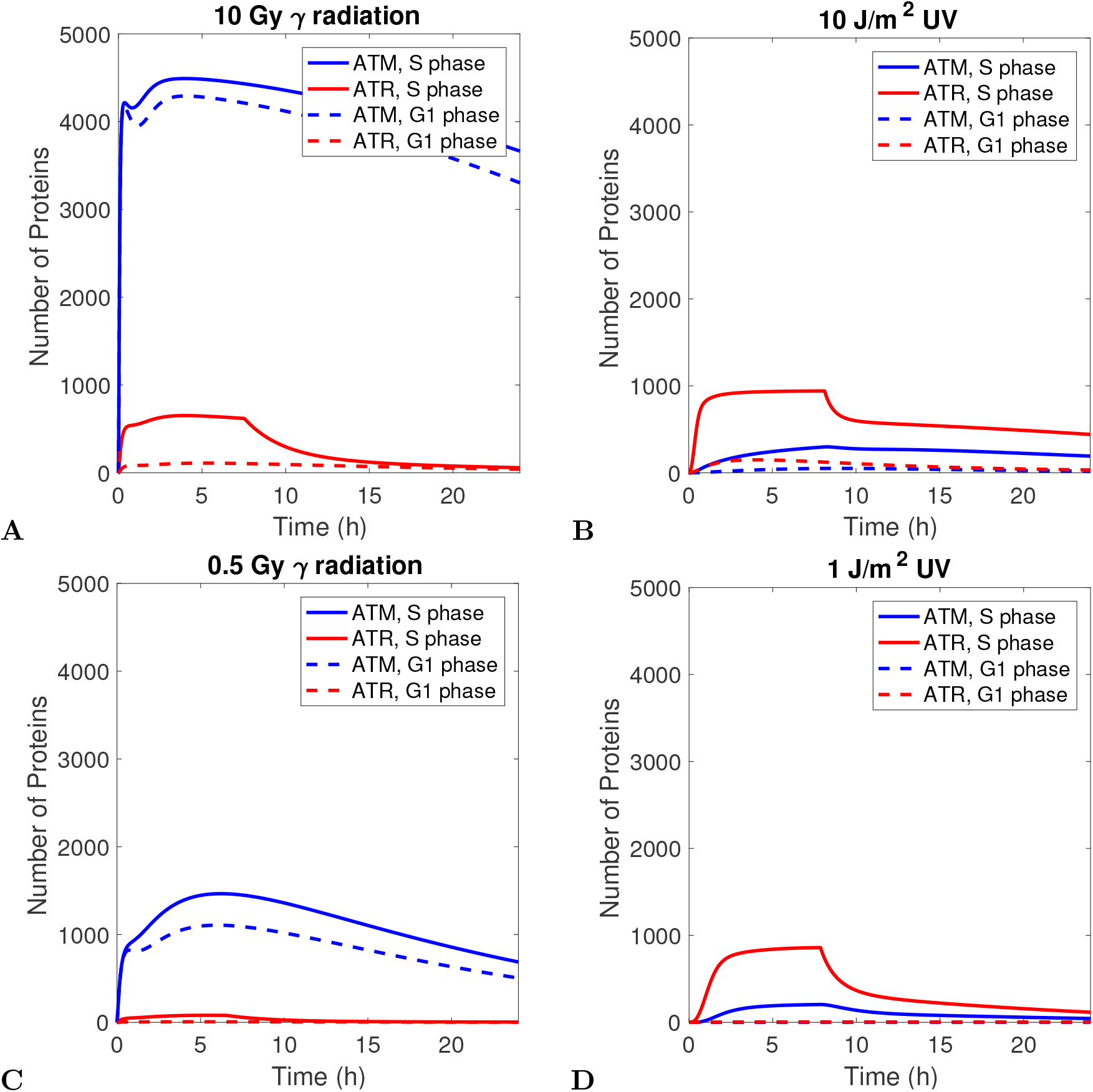
Comparison of ATM and ATR activity in response to high and low levels of damage: **(A)** 10 Gy *γ* radiation, **(B)**10 J/m^2^ UV, **(C)**0.5 Gy *γ* radiation, **(D)**1 J/m^2^ UV. In **(D)**, the lines representing ATM and ATR levels in G1 phase are nearly identical.

**Stiff et al. 2006 [19]:** This is an early work exploring how ATR activates ATM in response to UV damage. Because ATM phosphorylation does not require *γ*-H2AX, the authors conclude this activation is not primarily driven by spatial effects or repair intermediates. They instead argue that ATM Ser1981 is a substrate of ATR, which they support by performing a kinase assay with ATR and a Ser1981-containing peptide, and which later researchers confirm by substrate analysis [28].

**Bakkenist et al. 2003 [31]:** This work also reports on early ATM activation kinetics in response to small amounts of radiation. Bakkenist et al. report that exposure to as little as 0.4 Gy *γ* radiation is sufficient to provoke a robust ATM response within 15 minutes. As shown in Fig. 8C, even low levels of *γ* induce a fast response from the model. We present this with a caveat: 0.5 *γ* radiation to produces 2 C-DSBs on average, which is too few to model reliably with a continuous approximation. Variation in the number of complex lesions may strongly influence the dynamics at low radiation exposure.

The authors were unable to detect active ATM 1 hour after 10 J/m^2^ UV damage but do detect active ATM after 5 hours in S phase. Figure 8B shows ATM increasing gradually in S phase and reaching capacity after about 20 hours. We assume there is a certain threshold under which active ATM cannot be detected by Western blotting, and set this threshold at 100 proteins.

**Kousholt et al. 2012 [117]:** Kousholt et al. performed a series of experiments on the relationship between ATR signaling and DSB end resection. In cells with silenced CtIP, a protein that promotes end resection by MRN, Chk1 was initially activated at a rate comparable to that of the wild type, but its activity was not sustained. For a CtIP-deficient system (*k_MRN_* = 0), the model indeed predicts no change in initial ATR activation and lower levels of active ATR compared to the wild type (Figure 9A). The authors also show that ATR activates Chk1 before detectable end resection occurs. Our model attributes early activation of ATR to an ATM-dependent kinase cascade that likely involves TopBP1. Myers et al. have also published data that support these claims [131].

**Figure 9:**
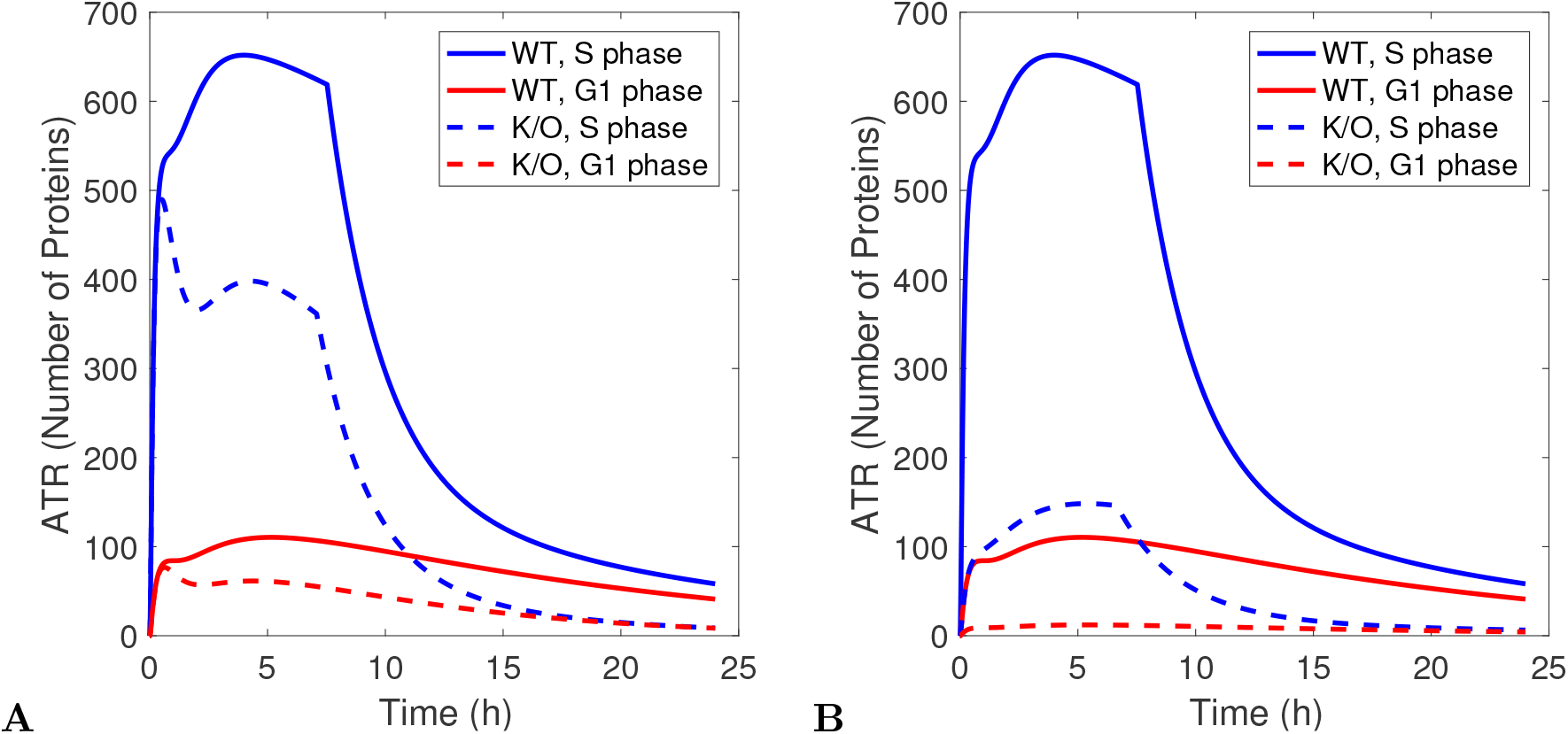
Post-*γ*-radiation ATR activity in the absence of certain key processes. **(A)** ATR activation in an end-resection-deficient cell, simulating the CtIP silencing experiment performed in [117]. **(B)** ATR activation in an ATR-kinase-suppressed cell, as in [20].

**Gamper et al. 2013 [20]:** Gamper et al. confirm that ATR can be activated outside of S phase and before the DSB ends undergo end resection. When the authors administered an ATR kinase inhibitor during G1 phase, ATR foci did not form around DSBs, which suggests the formation of these foci depends on some unidentified phosphorylation target of ATR. Since ATR kinase activity is required for active ATR to dissociate from RPA-bound ssDNA and allow inactive ATR to bind, we model ATR kinase deficiency, by lowering *k_AT R_* by an order of magnitude [132]. This abrogated ATR focus formation around DSBs in G1 phase (Fig. 9).

**Ward et al. 2004 [33]:** This early work shows that Chk1, an ATR substrate, is not upregulated during G1 phase in response to UV. It does not consider the ATR response to *γ* radiation, so it is not in conflict with later works claiming that ATR responds to DSBs in G1 phase. Since UV damage can induce a low amount of active ATR in G1 phase, (Fig. 8B), we assume there is a threshold under which active ATR cannot robustly activate Chk1, and estimate this threshold at 100 ATR molecules.

Table 6 summarizes the claims outlined in this section.

**Table 6:**
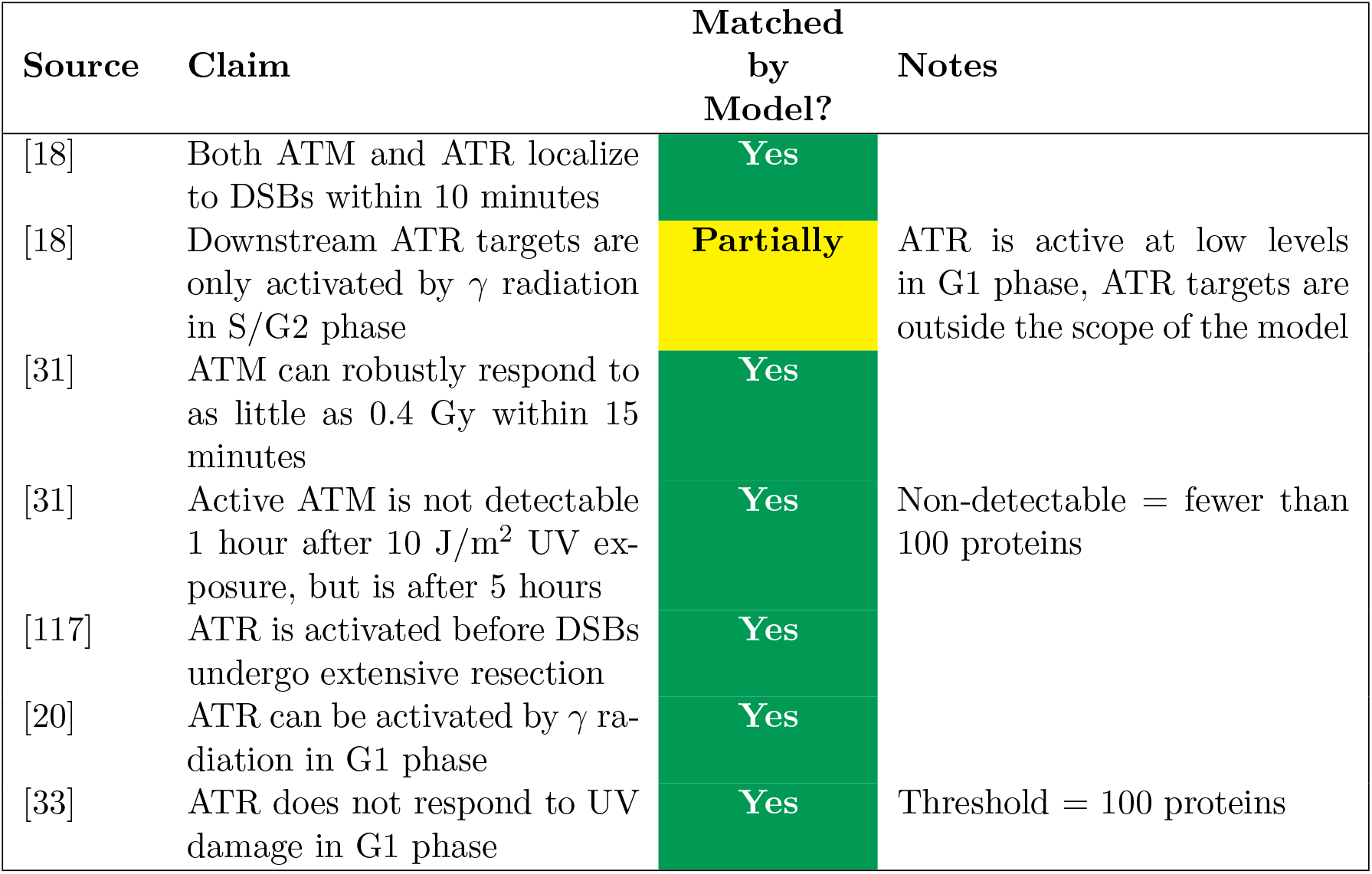
Comparison of the model results to crosstalk data, part 2.

**Tomimatsu et al. 2009 [124]:** These authors record the post-*γ*-radiation kinetics of ATR in cells with defective ATM (A-T cells). ATM/ATR substrates are still activated in these cells, which the authors attribute to ATR and DNA-PK. In contrast to the results of Jazayeri et al., ATR can colocalize to DSBs in cells with defective ATM, which the authors suggest is likely due to a redundant end resection pathway.

Figure 10B shows a simulation of the ATM knockout experiment described in Jazayeri et al. 2006 and Tomimatsu et al. 2009. In the wild-type case, ATR responds robustly to *γ* radiation within minutes, consistent with the observations of Jazayeri et al. When the cell is irradiated in G1 phase, ATR responds at a slower rate and with a lower carrying capacity compared to its activity in S phase. Knocking out ATM suppresses the initial speed, but not the carrying capacity, of ATR binding. Overall, the model’s behavior is consistent with the observation of Tomimatsu et al. that ATR responds to *γ* radiation in ATM-deficient cells.

**Figure 10:**
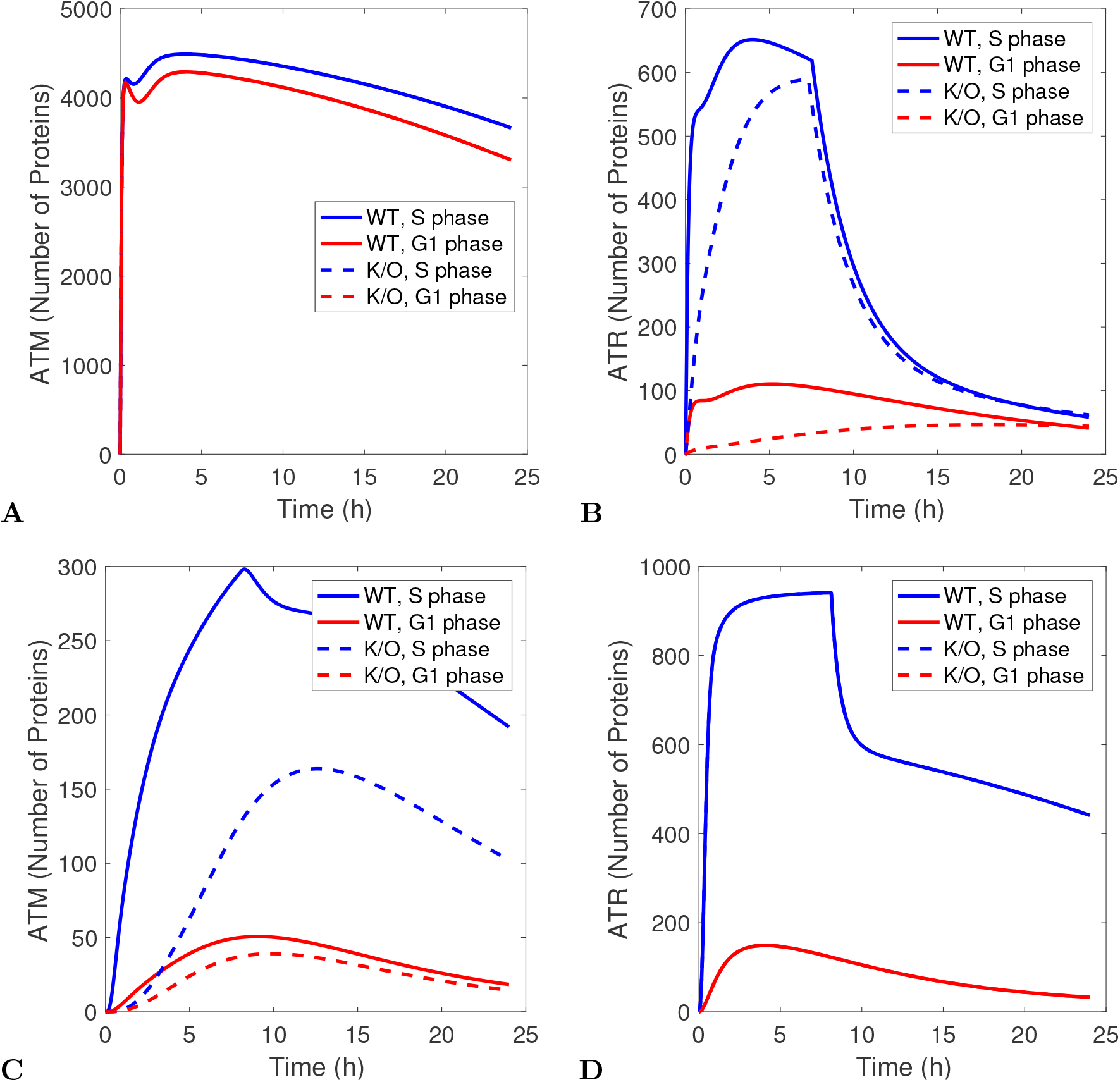
Predicted kinase activity in ATM or ATR knockouts. Results are for S and G1 phase. **(A)** ATM activation in response to *γ* radiation in an ATR K/O cell. **(B)** ATR activation in response to *γ* radiation in an ATM K/O cell. **(C)** ATM activation in response to UV radiation in an ATR K/O cell. **(D)** ATR activation in response to UV radiation in an ATM K/O cell. In **A** and **D**, the blue lines and the blue dashed lines are so close as to be indistinguishable, as are the red lines and the red dashed lines.

**Ray et al. 2016 [94]:** Ray et al. performed many of their experiments by exposing cells to 100 J/m^2^ UV, outside the dose range in which our model is valid. We only consider these authors’ experiments in which they exposed cells to 20 J/m^2^ UV. NER-deficient cells proficiently responded to UV damage during S phase, but formed fewer ATM/ATR foci after UV radiation in G1 phase.

Figure 11 shows the model’s predicted kinase activity in NER-deficient cells (*k_DN_* = *k_DT_* = 0). Removing NER did not preven the cell from responding to UV radiation in S phase, but completely suppressed both ATR and ATM during G1 phase. Our decision to change *k_DN_* and *k_DT_*, rather than *k_NER_*, reflects that the xeroderma pigmentosum (XP) cells used by Ray et al. have mutations that inhibit ssDNA creation and RPA-ATR interactions but do not necessarily slow repair after ATR is bound. The results shown are more consistent with XP-A and XP-C cells; to model XP-E cells, which are defective in GG-NER but proficient in TC-NER, we would set only *k_DN_* = 0.

**Figure 11:**
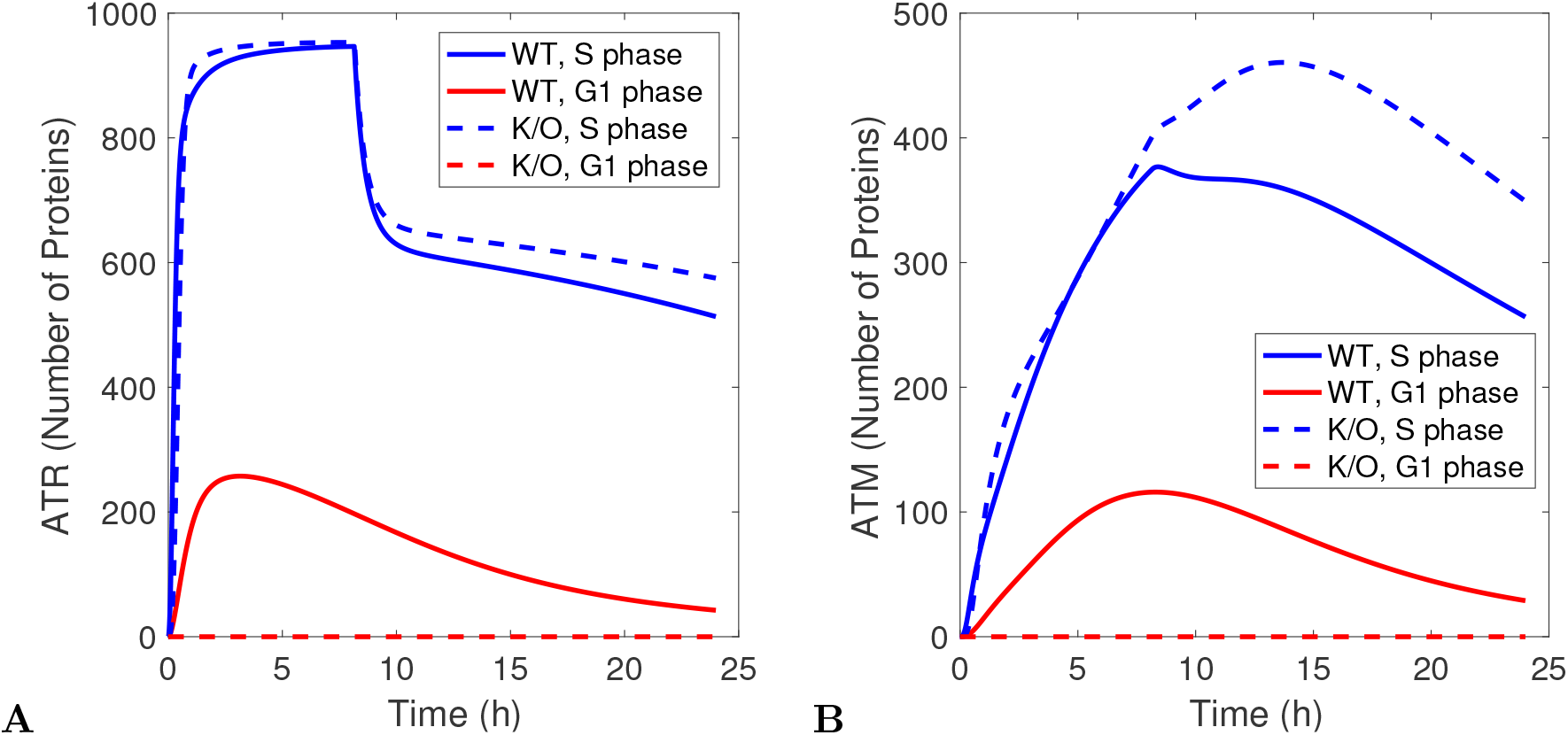
Model of the signal produced by **(A)** ATR and **(B)** ATM in WT and NER-deficient cells exposed to 20 J/^2^ UV. In both plots, the blue solid line (WT S phase activity) and blue dashed line (NER-deficient S phase activity) are so close as to appear indistinguishable.

**Mladenov et al. 2019-1 [133]:** This recent work focuses on how the activity of three PI3Ks— ATM, ATR, and DNA-PKcs—changes at high and low doses of *γ* radiation. At low doses of *γ* radiation, ATM and ATR cooperate to activate the G2 checkpoint, but at high doses of *γ* radiation, the cell can activate the G2 checkpoint when treated with either ATRi or ATMi. We cannot test the model against these claims because cell cycle arrest is beyond the scope of the model. We instead test two other results: that inhibiting ATR does not interfere with ATM activation during G2 phase, and that inhibiting ATM or ATR reduces the rate at which DSBs are end-resected.

As demonstrated in Fig. 8, the ATM response to *γ* radiation is not sensitive to cell cycle effects. Mimicking G2 phase by setting *s* = 1 and *G*(0) = 0 (not shown) produces a response similar to Fig. 10A and is similarly preserved when ATR is silenced. However, there are no mechanisms in the model that would allow ATR to affect end resection.

**Mladenov et al. 2019-2 [134]:** As a follow-up study, the same authors present experiments where the cell is radiated in S phase instead of G2. Knocking out ATM in S phase did not affect post-*γ*-radiation ATR signaling. Cells with end resection defects were proficient in ATR signaling, although, in contrast to the authors’ previous work, ATR had no effect on end resection during S phase. These results are consistent with previous experiments, and the simulations corresponding to these experiments appear in Figs. 10B and 9A.

Table 7 summarizes the claims outlined in this section.

**Table 7:**
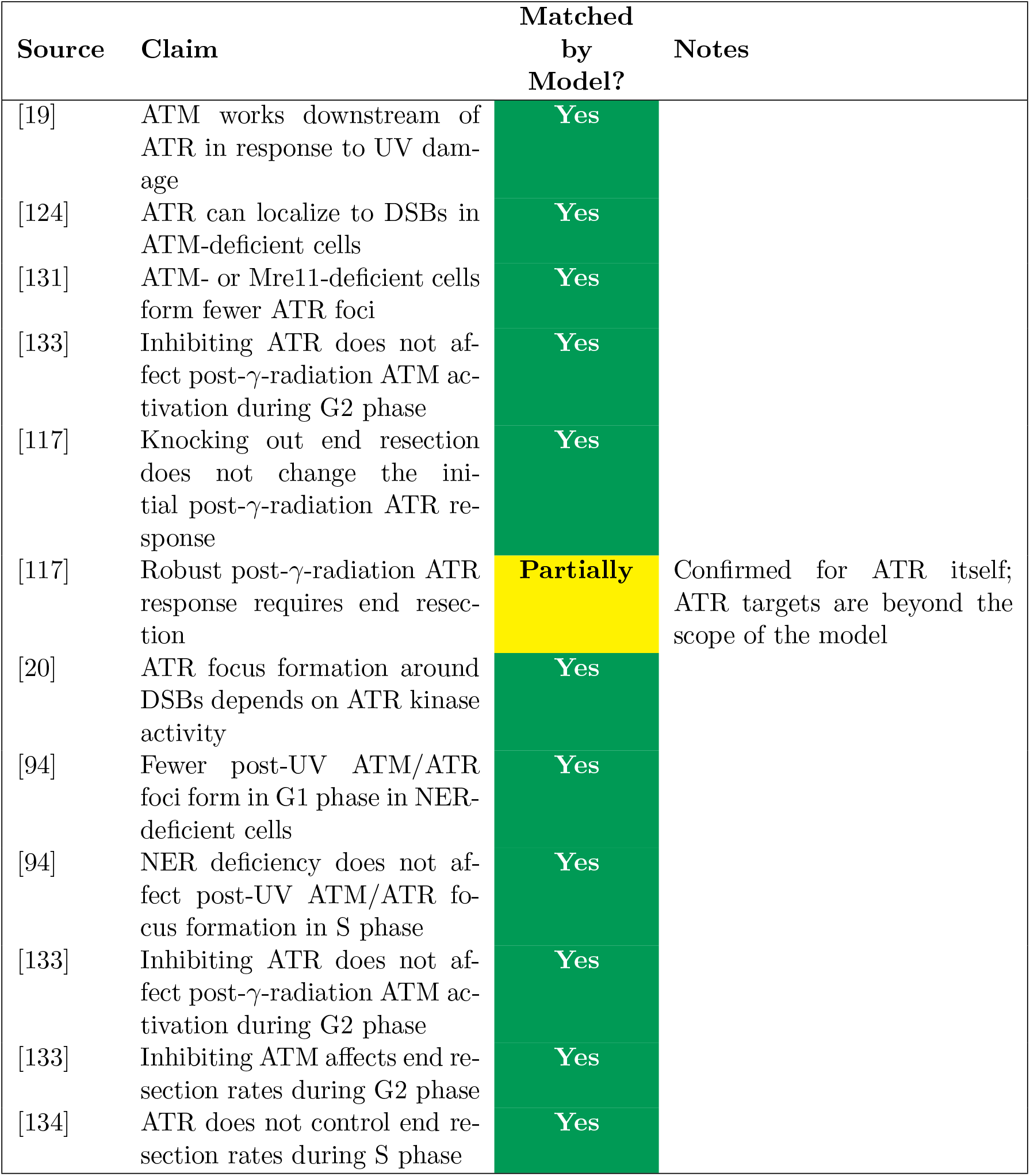
Comparison of the model results to crosstalk data, part 3.

## 4.9.4 Exceptions

Despite the number of interactions we consolidate, we have left out many elements of the ATM and ATR pathways. We list some notable results that are beyond the scope of the model.

**Batchelor et al. 2008 and 2011 [128, 22]:** ATM famously oscillates in response to *γ* radiation and other DSB-causing damaging agents, which causes similar oscillations in active p53. The mechanism that causes ATM to oscillate, dephosphorylation by Wip1, is p53-dependent. Because this model does not include p53, we do not expect ATM to oscillate in response to *γ* radiation.

**Vrouwe et al. 2011 [16]:** This 2011 work by Vrouwe et al. focuses on ATR activation in cells with defective NER. Cells with defective NER show a high, but delayed, accumulation of downstream ATR substrates. The authors speculate that, despite the NER deficiency, unrepaired photoproducts accumulate enough single-strand breaks to create ssDNA. While our model does not account for the side effects of strand breakage in repair-deficient cells, we do use their results suggesting that ATR is upregulated by NER outside of S phase.

We summarize the exceptions in Table 8.

**Table 8:** Comparison of the model results to crosstalk data, part 5.

| Source | Claim | Matched by Model? | Notes |
| --- | --- | --- | --- |
| [133] | Inhibiting ATR affects end resection rates during G2 phase | No | We do not include downstream effects of ATR on end resection |
| [128, 22] | Wip1 downregulation causes ATM to oscillate in response to $\gamma$ radiation | No | We do not include p53-dependent feedback loops |
| [16] | NER-deficient cells arrest in G1 phase after UV damage | No | We do not model ssDNA formation in repair-deficient cells |

## 4.10 Experimental conditions

We add or remove several key interactions to assess how they change the system’s qualitative behavior. The following subsections describe which components change to model these cases. Unless specified otherwise, the equations and parameters in the experimental system are the same as in the original model.

### 4.10.1 ATM kinase activity does not activate ATR

In the full model, both ATM and ATR promote ATR binding by locally activating TopBP1. We allow ATR to upregulate itself through TopBP1, but not ATM, by splitting each DSB class into three categories: the existing kinase-unbound and ATM- or ATR-bound categories, and an additional category for breaks bound to ATR. For example, we split DSBs into *L_DSB_*, *L_DSBA_*, and *L_DSB,AT R_*:

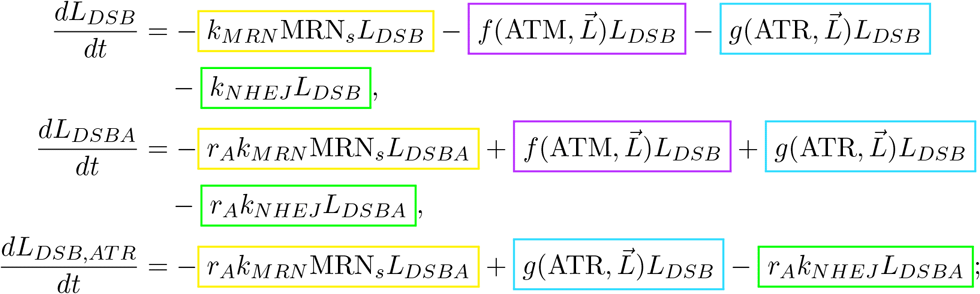

with associated ATR class

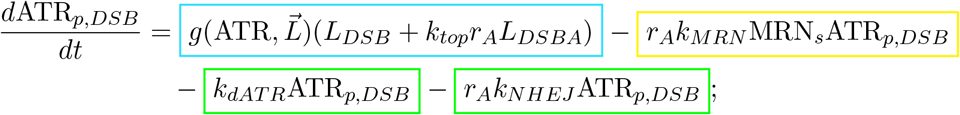

and with

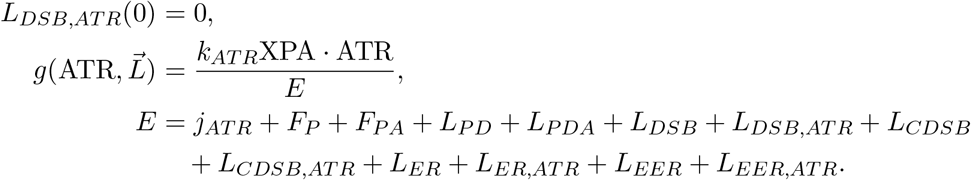

The ATM classes do not change. We label the interactions related to this experimental condition in Fig. 3.

### 4.10.2 ATR kinase activity does not activate ATM

ATR directly binds to and activates ATM *k_aa_* times as fast as DSBs activate ATM. We set *k_aa_* = 0 to remove this interaction and indicate this change in Fig. 5.

### 4.10.3 ATM and ATR do not phosphorylate H2AX

We assume histones are phosphorylated quickly after either kinase binds, and all subsequent protein interactions with the damaged site are *r_A_* times faster. To remove the effects of histone phosphorylation, we set *r_A_* = 1. We label the interactions affected by this change in Fig. 2.

### 4.10.4 Replication stress on MDS activates ATR

Gamma radiation induces many non-DSB lesions, some of which can cause replication stress during S phase. To account for post-*γ*-radiation replication fork stalling, we use MDS as a representative class of persistent non-DSB lesions. We partition unprocessed MDS into non-replicated and replicated DNA classes (*L_MDS_*, *L_MDSR_*), where only lesions in *L_MDS_* can cause replication stress; and include forks stalled on MDS in ATR-unbound and -bound states (*F_s,MDS_*, *F_s,MDSA_*). These equations govern the four MDS classes:

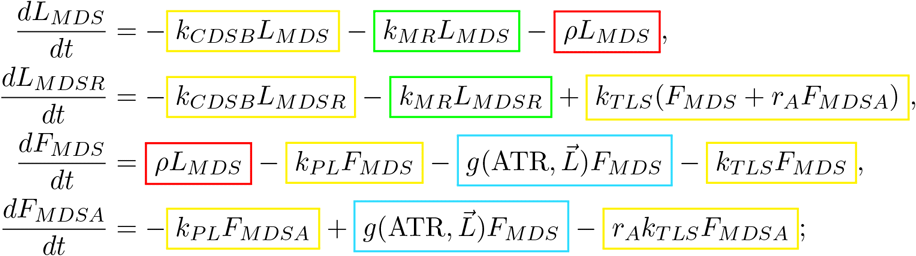

with associated kinase classes

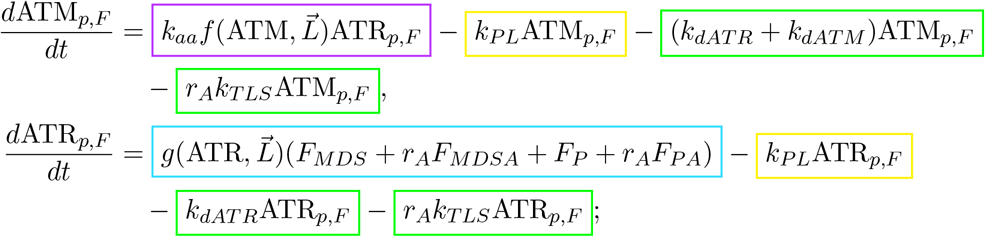

and

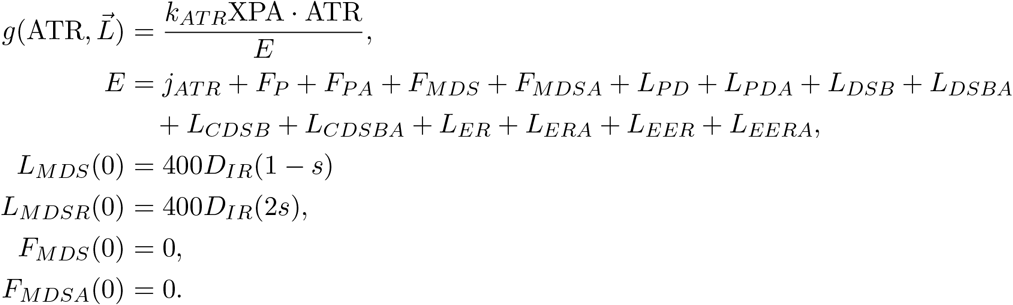

In particular, if TLS repairs all replication forks at the same rate, we do not need separate classes for ATR bound to forks stalled on photoproducts and MDS. All four new classes break to form DSBs, where DSBs created by endemic breakage of stalled replication forks move into the end-resected class:

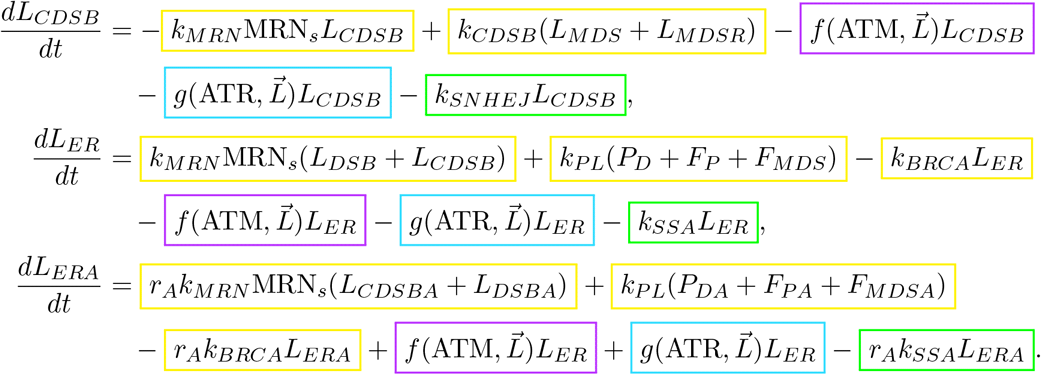

This case adds one new parameter: *k_MR_*, the repair rate of MDS, which we set to 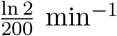. We label the class of lesions added to accommodate to this experimental condition in Fig. 1.

### 4.10.5 ssDNA breaks to form DSBs

The parameter *k_pL_* controls the rate at which ssDNA endemically breaks to form DSBs. We set *k_pL_* << 1 for ssDNA breakage and label the state transitions affected by this change in Fig. 5.

### 4.10.6 ATM dissociates from end-resected DSBs

Based on the hypothesis set by Shiotani and Zhou that ATM dissociates from DSBs upon end resection, we remove ATM_*p,ER*_ and ATM_*p,EER*_ the end-resected-DSB-bound ATM classes [116]. In addition, neither end-resected DSB class can move to the kinase-bound class by interacting with ATM:

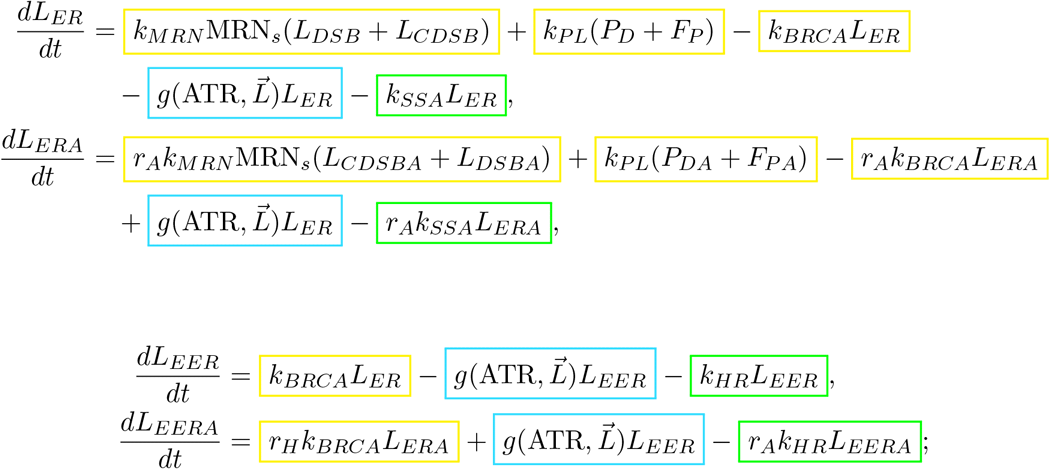

and we change the ATM activation term to

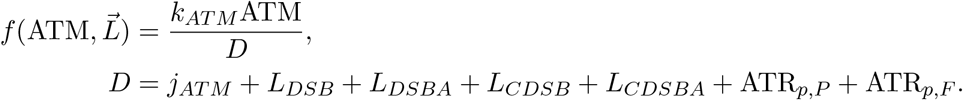

We label the ATM classes affected by this experimental condition in Figs. 3 and 5.

### 4.10.7 ATR dissociates from extensively-end-resected DSBs

Rad51 displaces RPA on ssDNA to initiate extensive end resection. To account for the possibility that RPA does not re-bind to the ssDNA after being outcompeted, we remove ATR_*p,EER*_, the extensively-end-resected ATR class, prevent extensively end-resected DSBs from binding to ATR:

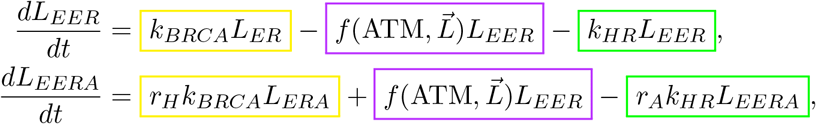

and change the ATR activation term to

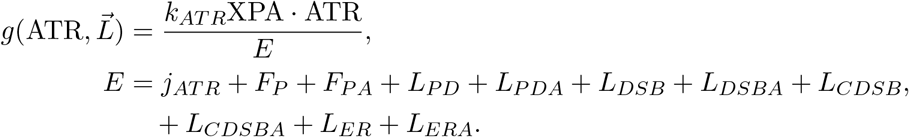

We label the ATR class affected by this experimental condition in Figs. 3 and 5.

## 5 Results

The ATM and ATR signaling dynamics predicted by the base model are consistent with over 20 experimental observations laid out in Section 4.9. To test how modes of crosstalk influence ATM and ATR signaling, we add or remove seven activation mechanisms as outlined in Section 4.10. Table 9 indicates whether the altered model is consistent with those same experimental observations. When inconsistencies arise, we briefly discuss what they imply about damage response signaling. Lastly, we introduce the cell to radiation at different times to the cell cycle relative to the start of S phase and record the change in ATM and ATR response.

**Table 9:**
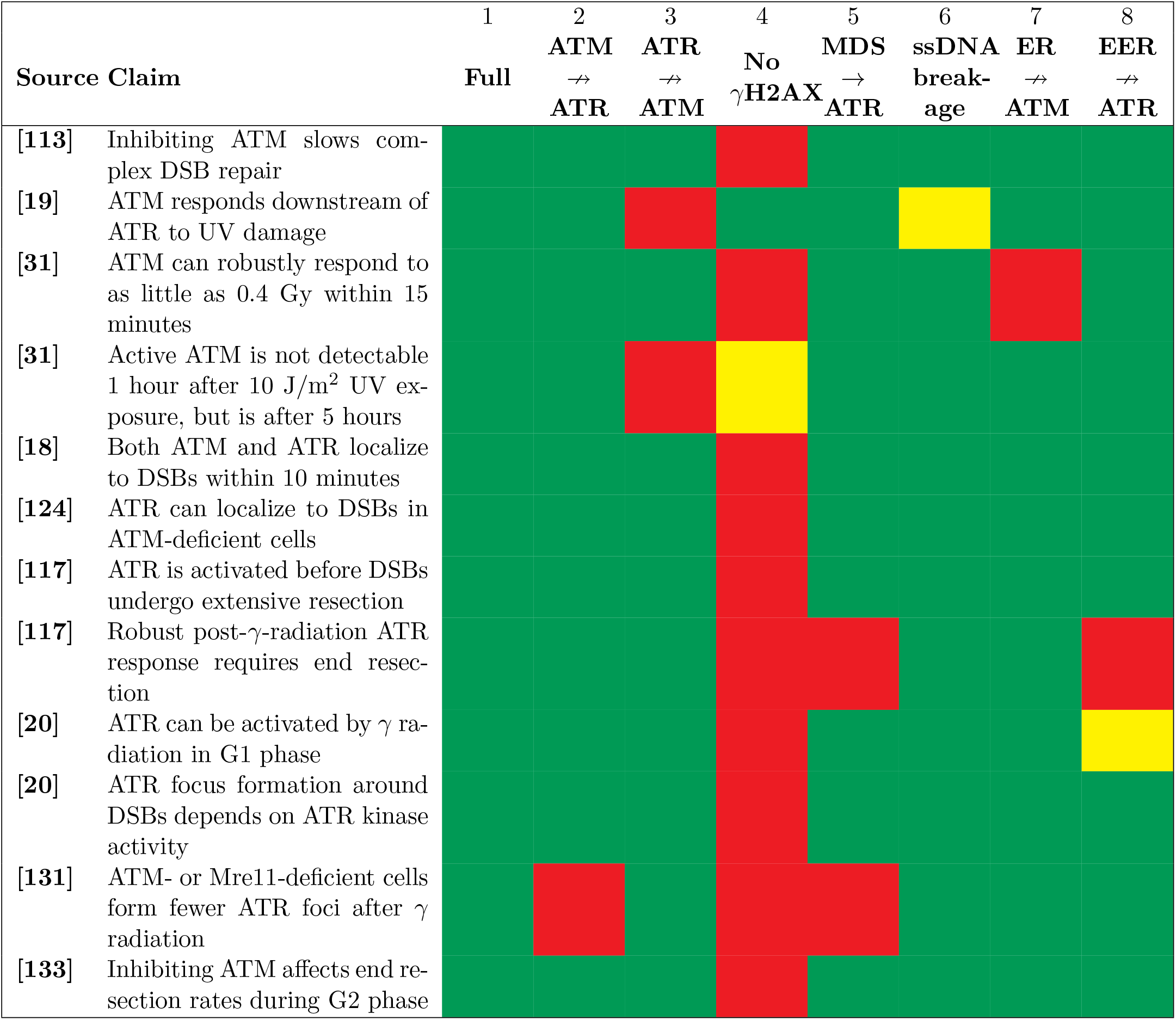
Changes in the qualitative model validation with or without key processes. Some rows are missing because all eight models satisfied the claims they represent. **Column 1:** the original model. **Column 2:** Model without ATM kinase activation of ATR. **Column 3:** Model with *k_aa_* = 0, removing ATR kinase activation of ATM. **Column 4:** Model with *r_H_* = 1, removing increase in binding rates to kinase-bound lesions. **Column 5:** Model with replication fork stalling on *γ*-radiation-induced lesions. **Column 6:** Model with *k_P L_* = 1×10^*−*7^, adding the possibility that ssDNA can endogenously break to form DSBs. **Column 7**: Model with ATM_*p,ER*_ and ATM_*p,EER*_ removed to represent the hypothesis that ATM dissociates from end-resected DSBs [116]. **Column 8**: Model with ATR_*p,EER*_ removed to represent the possibility that RPA cannot re-bind to Rad51-bound ssDNA after being displaced.

### 5.1 Wild-type model dynamics

Figure 12 shows the time evolution of both active kinases in response to *γ* or UV radiation administered at the start of S phase. Although the time course is unrealistic for an actively replicating cell, we show how the signal would evolve over 96 hours if downstream regulatory pathways did not suppress ATM or ATR activation. This simulated cell has 5000 ATM molecules and 1000 ATR molecules, reflecting the result in Wang et al. that most human organs have more ATM than ATR [130].

**Figure 12:**
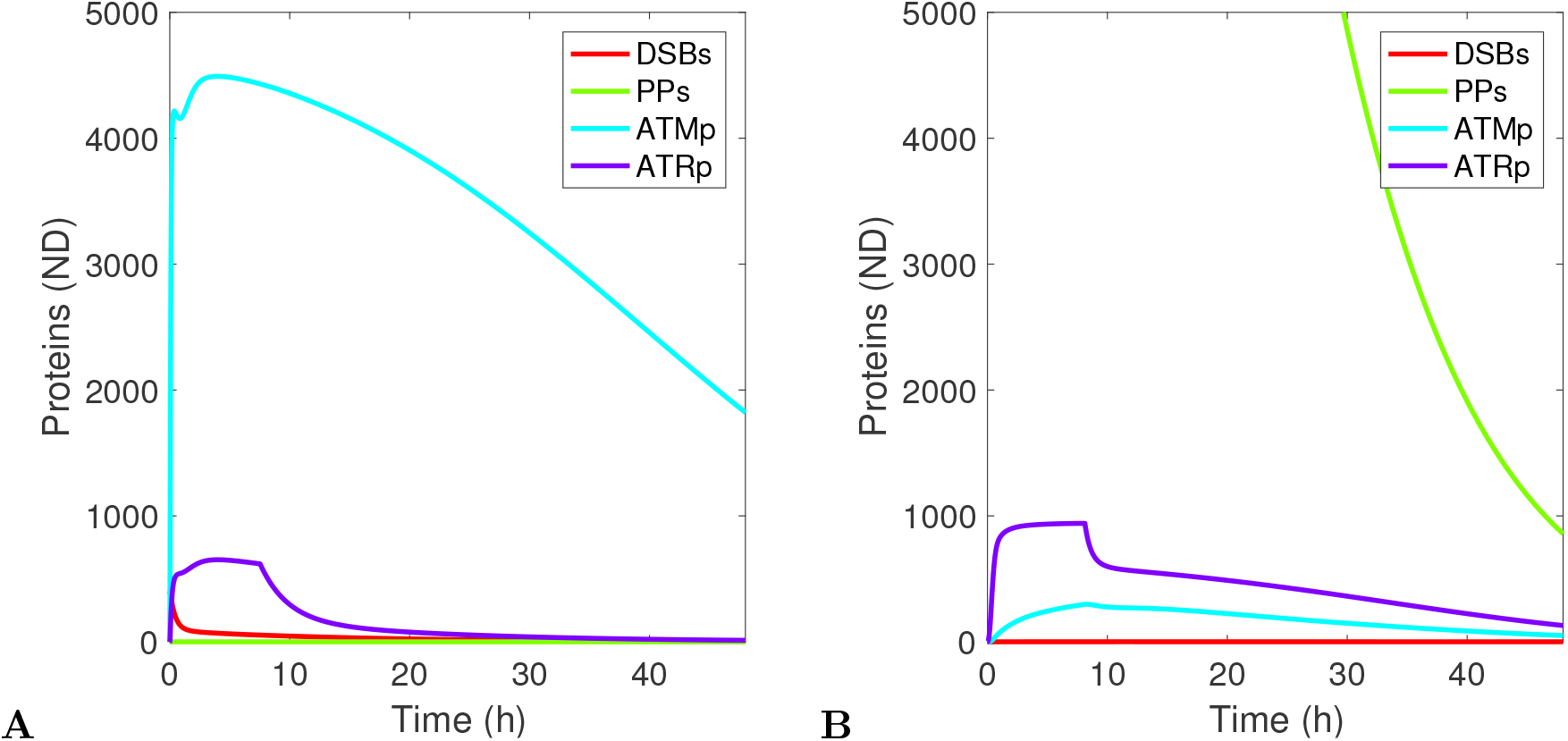
Representative behavior of both kinases for 48 hours after exposure to **(A)**10 Gy *γ* radiation or **(B)**10 J/m^2^ UV at the beginning of S phase. In **B**, the initial number of photoproducts induced by 10 J/m^2^ UV radiation is 45,000.

After *γ* irradiation, almost every ATM and ATR molecule in the cell becomes active within an hour of exposure. Active ATR declines two hours after irradiation as simple DSBs are repaired, and continues to decline over the course of three days as fewer persistent breaks remain. ATM remains active even when only a few complex DSBs remain in the cell. This reflects observations that only one or two DSBs can effectively activate ATM, although stochastic effects may dominate in a system with few lesions.

ATR responds within minutes of the start of S phase in a UV-irradiated cell and persists at a high-activity level that remains constant over 96 hours. ATM rises slowly, reaching a maximum almost a day after exposure. Although 1 J/m^2^ UV radiation produces thousands of photoproducts and 1 Gy *γ* radiation produces only four complex DSBs, the combined response of ATM and ATR to UV damage is lower.

### 5.2 ATM and ATR mutual upregulation drives early kinase dynamics

ATM interacts with DSBs and is canonically associated with *γ* radiation signaling, while ATR interacts with RPA-bound ssDNA and is canonically associated with UV radiation and replication stress signaling. Each kinase can respond to its non-canonical form of damage through one of three mechanisms: ATR operates downstream of ATM and vice versa, ATM and ATR modify the lesion to encourage interactions with regulatory proteins, and the lesion itself can undergo structural changes that interact with its non-canonical kinase. We individually removed each of these mechanisms to assess how they affect secondary kinase behavior.

ATM activates TopBP1, a mediator required for ATR binding. Column 2 of Table 9 describes the qualitative behavior of a model where ATM does not activate ATR through a kinase cascade. Although removing this interaction lowered the amount of active ATR following *γ* irradiation, the only claim this model violated was that fewer ATR foci develop in ATM-deficient cells.

Active ATR directly phosphorylates ATM. If we remove this interaction, as shown in Column 3 of Table 9, not only does ATM no longer depend on ATR to be activated after UV damage, it cannot act quickly enough to match experimental observations. Single-strand DNA breakage also activates ATM after UV damage but was not strong enough to restore ATM activity during the first five hours after irradiation.

In the third form of mutual upregulation, ATM and ATR can cause the lesion to undergo conformational changes, such as histone unwinding, that promote interaction with both kinases. As shown in Column 4 of Table 9, when we remove this interaction, the resulting model fails almost every claim. It even violated observations unrelated to crosstalk: ATM did not pass the detectability threshold in response to 10 Gy *γ* radiation. Despite the kinases responding weakly, their kinetic properties such as initial activation speed, time to maximal activation, and signal duration were similar to those of the original model.

### 5.3 Changes to lesion structure do not initiate secondary kinase activation

Gamma radiation induces RPA-bound ssDNA in one of three ways: RPA can bind to the 1-10 bp ssDNA tails on unprocessed DSBs, DSBs can undergo end resection to lengthen these ssDNA tails, and replication forks can stall on any radiation-induced lesion. The full model includes the first two processes. When we allow replication forks to stall on *γ*-radiation-induced lesions (Table 9, Column 5), replication stress drives ATR activation during S phase. This conflicts with experiments showing that active ATR primarily localizes to DSBs in all stages of the cell cycle. In particular, Kousholt et al. identify DSB end resection as the mechanism that prolongs ATR signaling after five hours, but replication fork stalling dominates ATR activation so thoroughly that suppressing end resection does not affect the S-phase signal [117].

DSBs can form after UV damage when ssDNA breaks. Adding a low rate of ssDNA breakage did not cause the model to violate any claims (Table 9, Column 6). If the ssDNA breakage rate is too high, ATM can activate independently of ATR after UV exposure. We found that *k_pL_* = 1 × 10^*−*7^ is a reasonable estimate for which all claims hold. Hence, aside from DSB end resection, changes in lesion structure due to cellular processing do not initially activate secondary kinases; however, it is still possible for them to influence long-term signaling behavior.

### 5.4 Interactions with end-resected DSBs lengthen kinase signals

As DSBs undergo end resection, they gain longer ssDNA tails and lose the short ends that initially attract ATM. In a model where end-resected DSBs were incapable of activating ATM (Table 9, Column 7), the cell cannot respond to low levels of ionizing radiation. Moreover, while ATM and ATR could both respond proficiently to 10 Gy *γ* radiation, both kinases decay to undetectable levels within 8 hours. This is at odds with experiments in which, with its negative regulator Wip1 silenced, ATM produces a strong signal 10 hours after exposure to 10 Gy *γ* radiation [128].

Rad51 drives extensive end-resection by outcompeting RPA on ssDNA. If RPA cannot re-bind to ssDNA after being outcompeted, extensively-end-resected DSBs would not be able to upregulate ATR. When we prevent this DSB class from activating ATR (Table 9, Column 8), the post-*γ*-radiation ATR signal is transient, lasting for only five hours. This model also predicts a stronger ATR response in end-resection-deficient cells, when removing end resection should instead attenuate long-term ATR activation.

### 5.5 The cell cycle influences ATM and ATR activation

To understand how the cell cycle affects ATM and ATR activity, we modeled their response to damage induced at different times relative to the start of S phase. Figure 13 shows the cell-cycle-specific activity of a cell with a 16-hour G1 phase and a 6-hour S phase in damage-free conditions. We induced damage during G1 phase or S phase at times separated by 1 hour relative to the start of S phase.

**Figure 13:**
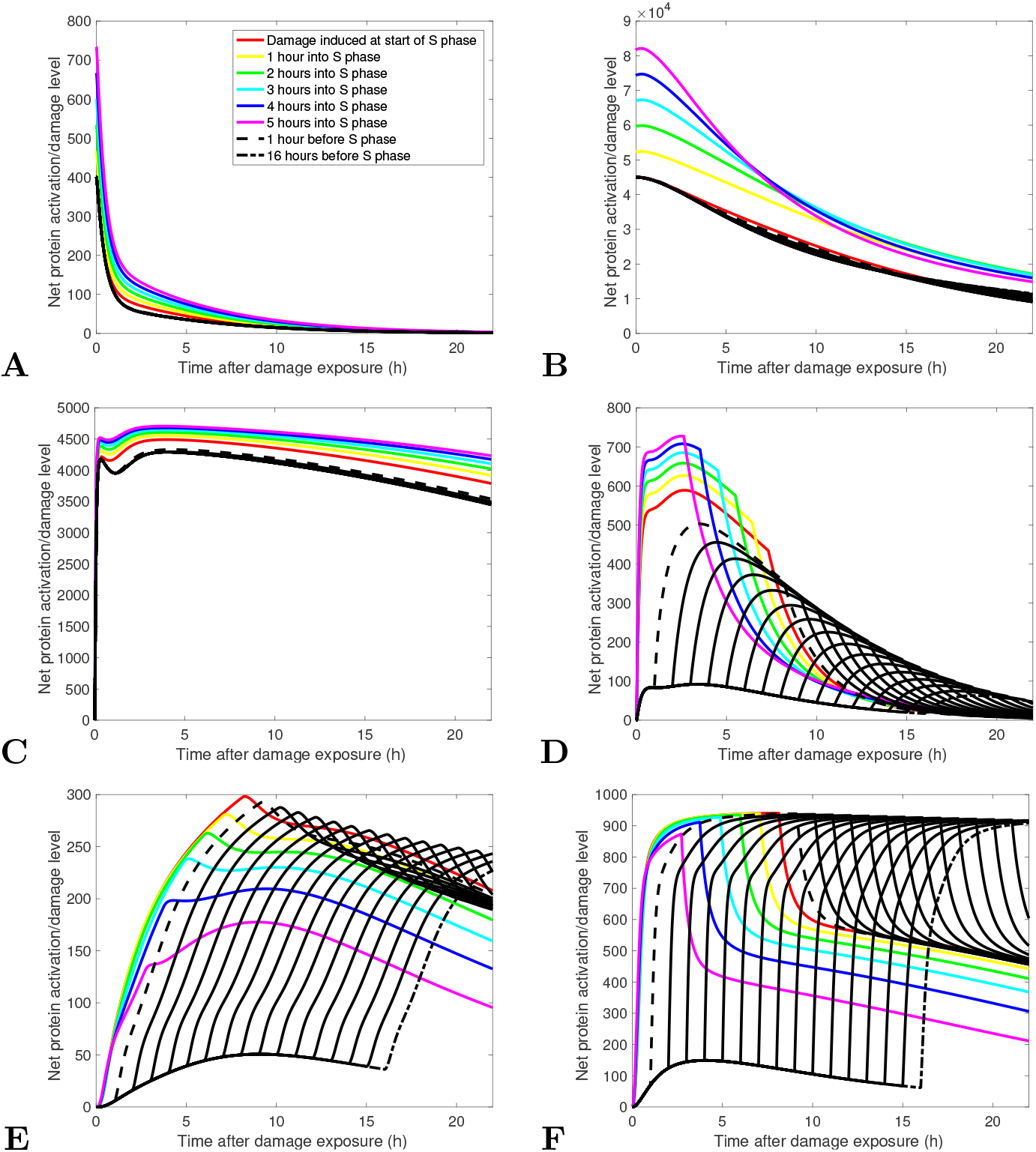
Effects of the cell cycle on damage response. Each line represents a cell damaged at a different time relative to the start of S phase, which we model as lasting 6 hours in a healthy cell. Colored lines represent cells damaged during S phase. Black lines represent cells damaged between 1 and 16 hours before the start of S phase, where signals progress monotonically. These diagrams show (**A**) DSB repair, (**B**) photoproduct repair, (**C**) ATM activity after *γ* irradiation, (**D**) ATR activity after *γ* irradiation, (**E**) ATM activity after UV irradiation, (**F**) ATR activity after UV irradiation.

Since photoproducts on stalled replication forks require an additional step to repair, the cell cycle affects photoproduct repair rates (Fig. 13B) but not DSB repair rates (Fig. 13A). This agrees with experimental observations. Radiation creates more lesions during S phase because it can react with the extra DNA synthesized during replication.

ATM is robustly activated by *γ* radiation in both G1 and S phase (Fig. 13C). ATR responds most strongly to *γ* radiation induced during S phase (Fig. 13D). If the cell is irradiated during G1 phase, ATR responds weakly to *γ* radiation until the start of S phase, when it increases sharply relative to the amount of remaining DSBs. This scenario would not happen in a real cell, since targets of active ATM would arrest the cell cycle in G1 phase until no breaks remain.

If we expose the cell to UV in G1 phase, or close to the start of S phase, ATR quickly reaches a maximum and equilibrates within 5 hours to a high level (Fig. 13E). ATM does not achieve the same maximum if the cell is damaged during S phase, but converges to a similar high-activity equilibrium, albeit at a reduced rate that lowers as the cell is damaged later in S phase. UV damage activates ATM most efficiently when it is administered close to the start of S phase (Fig. 13F). If we damage the cell in the middle of S phase, ATM operates downstream of a reduced amount of ATR, and if we damage the cell early in G1 phase, it has more time to repair photoproducts before they induce replication stress, lowering the number of stalled forks that endemically break. We do not model metaphase, and understand G2 phase to be the state of the model after it completes replication.

### 5.6 Endemic breakage enhances post-UV kinase signal variation

Figure 14 shows how the area under the active ATM or ATR curve varies with radiation dose over 24 hours. Increasing the *γ* radiation dose raises the level of both active ATM and active ATR with diminishing results (Fig. 14A). In a cell with ten times less ATR (Fig. 14C), the change in active ATM remains the same.

**Figure 14:**
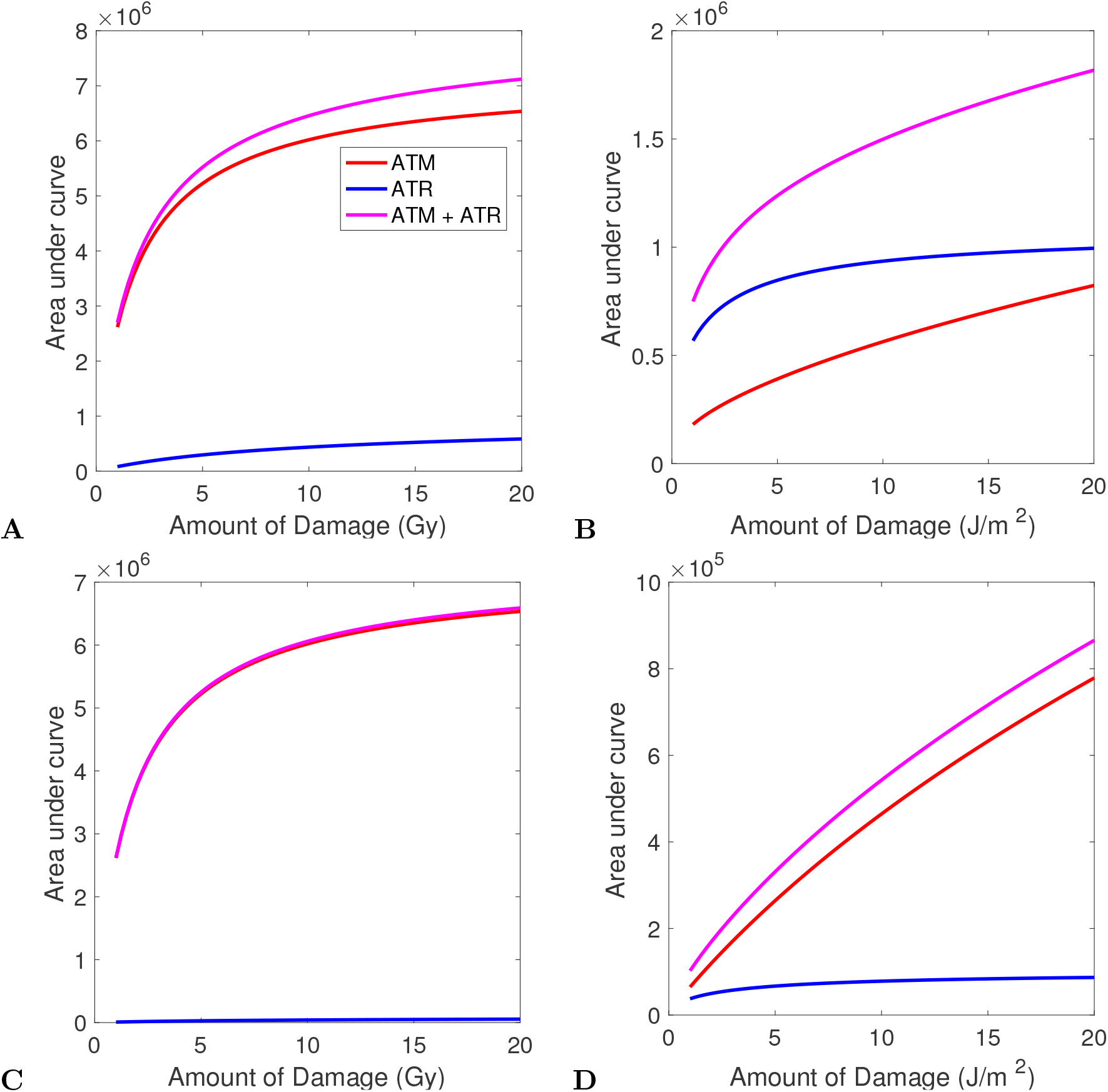
Dose response curves for ATM and ATR. **(A)** Response to varying doses of *γ* radiation; ATM_*tot*_ = 5000, ATR_*tot*_ = 1000. **(B)** Response to UV radiation; ATM_*tot*_ = 5000, ATR_*tot*_ = 1000. **(C)** Response to *γ* radiation; ATM_*tot*_ = 5000, ATR_*tot*_ = 100. **(D)** Response to UV radiation; ATM_*tot*_ = 5000, ATR_*tot*_ = 100.

The cell is sensitive enough to UV damage that mutual targets of ATM and ATR show graded responses to UV damage in the range of 2–10 J/m^2^ [128]. We therefore expect total kinase activity to vary over this range. As shown in Fig. 14B, in a cell with the same total amounts of ATM and ATR given in Table 4, UV damage activates ATR quickly and efficiently even at low levels of exposure. Both ATM and ATR increase over the given UV dose range. For a cell with ten times less ATR (Fig. 14D), ATR is not sensitive to changes in UV dose; it becomes maximally active even at low doses of UV radiation. Instead, active ATM levels are what cause the net signal to vary. This happens because more ssDNA breaks to form DSBs as the number of photoproducts in the cell increases.

## 6 Discussion

We introduce the first mathematical model of ATM and ATR co-activation, one that satisfies 21 claims from various experimental sources. This model includes three modes of ATM/ATR crosstalk: mutual upregulation of ATM and ATR, enhanced protein recruitment to lesions bound by ATM or ATR, and changes in lesion structure caused by cellular processing.

Altering these modes of crosstalk provides insight into how each mode affects ATM and ATR signaling. While mutual upregulation drives the activation of both kinases in the early stages of damage response, changes in lesion structure due to cellular processing are what prolong kinase activation in response to both *γ* and UV radiation.

The model also has cell-cycle-specific functionality for damage induced in S or G1 phase. Replication stress, preferential activation of ATR during S phase, and replication slowing by ATR all change the model’s predicted kinase activity during S phase. But while ATR is indispensable for promoting apoptosis in response to replication stress, if nearly all ATR molecules become active shortly after replication stress appears, secondary kinases such as ATM are what cause the combined pro-apoptotic signal to vary with damage intensity. Overall, this model successfully resolves qualitative experimental claims about ATM and ATR crosstalk that have been collected over the past 17 years.

### 6.1 Kinase dynamics

The three ATR and ATM crosstalk mechanisms operate on different timescales. Immediately after radiation exposure, ATM and ATR kinase activity drive their activation. Structural changes to the lesions prolong the combined kinase signal: in the model, double-stranded break end resection sustains ATM and ATR hours after *γ* irradiation, and ssDNA breakage activates ATM for days after exposure to UV light.

Although the model requires end-resected DSBs to drive kinase activation, experimentalists have shown that the long ssDNA tails on end-resected DSBs cannot bind to ATM. To reconcile these results, we suggest that ATM may bind to the breaks before end-resection and then drives its own activation through autophosphorylation; that is, without directly binding to the break. This supports a model in which DSB foci may not necessarily be held together by chemical bonds, but could also form as a cloud of active proteins around the original lesion.

Thousands of UV photoproducts do not constitute a lethal amount of damage, but a cell can apoptose in response to only a few sustained double-strand breaks. We do not yet understand how kinase signaling distinguishes between small amounts of lethal damage and large amounts of non-lethal damage. In this model, *γ* radiation activates almost all ATM and ATR molecules in the cell within minutes, and the signal it produces can be sustained by a single double-strand break. UV damage effectively activates ATR and produces an ATM signal that reaches a maximum hours after irradiation.

The effect on UV-induced single-strand DNA breakage on ATM activation also has fascinating implications. A recent proteomics study suggests that ATM is much more abundant in human cells than ATR. In two organs, the heart and esophagus, the researchers detected no ATR. We considered whether cells with low ATR levels could distinguish between high and low dises of UV. In these cells, UV radiation produces an abundance of non-lethal photoproducts that cause high ATR activity regardless of the extent of damage. Because ATR is so thoroughly activated by even small numbers of photoproducts, it seems incapable of distinguishing between small and large doses of UV damage. Instead, larger doses of UV damage cause repair processes to create more singlestrand DNA. This increases the number of single-strand DNA regions that break endemically and activate ATM. In cells with naturally low levels of ATR, we therefore expect ATM, not ATR, to produce a signal that varies with the strength of UV doses.

### 6.2 Unresolved issues

Spatial properties affect several model components. Chromatin unwinding, DSB focus geometry, and diffusion of regulatory proteins away from the break can only be addressed indirectly in a non-spatial model. In particular, DSB foci may attract more regulatory proteins as they grow, but our model treats increased interaction rates like a switch, which turns on fully when the break has interacted with at least one PIKK-family kinase. Modeling either the three-dimensional focus size or the number of regulatory molecules in the focus would be more realistic, and would also be more mathematically challenging to construct. Lastly, because prolonged signals in response to *γ* radiation require few complex DSBs, stochastic interactions become more important in the late stage of the cell’s ionizing radiation response.

The experimental papers we use to validate the model represent different cell lines, methodology, and time scales. Often, instead of recording changes in active ATR over time, authors infer ATR activation from its downstream effects, such as Chk1 activation or the prevalence of G2 arrest. This led in some cases to different papers presenting seemingly contradictory claims about ATR activity. Because we base our parameter estimates on data sets across a range of cell types, they do not attempt to quantitatively predict cellular behavior.

Downstream regulatory pathways cause ATM to oscillate in response to *γ* radiation. In a future work, we will present a continuation of this model that incorporates ATM auto-suppression as part of p53 autoregulatory function.

This model highlights the importance of treating DNA damage signaling as a holistic effort rather than attributing it to a single dominant kinase. Our model suggests that ATM and ATR each convey crucial information to the cell’s regulatory pathways in response to both *γ* and UV radiation. One particular substrate of ATM and ATR, p53, is a damage aggregator responsible for interpreting a wide array of stressors. To understand how p53 makes decisions about cell fate, we must understand the total signal p53 receives from its upstream damage detection pathways. ATM and ATR are two of many kinases that contribute to this total signal. We hope these results inspire other researchers to seek a deeper understanding of how crosstalk between DNA damage signaling kinases drives decisions about cell fate.

## Notes

### Competing Interest Statement

The authors have declared no competing interest.

